# Engineering the thermotolerant industrial yeast *Kluyveromyces marxianus* for anaerobic growth

**DOI:** 10.1101/2021.01.07.425723

**Authors:** Wijbrand J. C. Dekker, Raúl A. Ortiz-Merino, Astrid Kaljouw, Julius Battjes, Frank W. Wiering, Christiaan Mooiman, Pilar de la Torre, Jack T. Pronk

## Abstract

Current large-scale, anaerobic industrial processes for ethanol production from renewable carbohydrates predominantly rely on the mesophilic yeast *Saccharomyces cerevisiae*. Use of thermotolerant, facultatively fermentative yeasts such as *Kluyveromyces marxianus* could confer significant economic benefits. However, in contrast to *S. cerevisiae*, these yeasts cannot grow in the absence of oxygen. Response of *K. marxianus* and *S. cerevisiae* to different oxygen-limitation regimes were analyzed in chemostats. Genome and transcriptome analysis, physiological responses to sterol supplementation and sterol-uptake measurements identified absence of a functional sterol-uptake mechanism as a key factor underlying the oxygen requirement of *K. marxianus*. Heterologous expression of a squalene-tetrahymanol cyclase enabled oxygen-independent synthesis of the sterol surrogate tetrahymanol in *K. marxianus*. After a brief adaptation under oxygen-limited conditions, tetrahymanol-expressing *K. marxianus* strains grew anaerobically on glucose at temperatures of up to 45 °C. These results open up new directions in the development of thermotolerant yeast strains for anaerobic industrial applications.

---

In terms of product volume (87 Mton y^−1^)^1,2^, anaerobic conversion of carbohydrates into ethanol by the yeast *Saccharomyces cerevisiae* is the single largest process in industrial biotechnology. For fermentation products such as ethanol, anaerobic process conditions are required to maximize product yields and to minimize both cooling costs and complexity of bioreactors^3^. While *S. cerevisiae* is applied in many large-scale processes and is readily accessible to modern genome-editing techniques^4,5^, several non-*Saccharomyces* yeasts have traits that are attractive for industrial application. In particular, the high maximum growth temperature of thermotolerant yeasts, such as *Kluyveromyces marxianus* (up to 50 °C as opposed to 39 °C for *S. cerevisiae*), could enable lower cooling costs^6–8^. Moreover, it could reduce the required dosage of fungal polysaccharide hydrolases during simultaneous saccharification and fermentation (SSF) processes^9,10^. However, as yet unidentified oxygen requirements hamper implementation of *K. marxianus* in large-scale anaerobic processes^11–13^.

In *S. cerevisiae*, fast anaerobic growth on synthetic media requires supplementation with a source of unsaturated fatty acids (UFA), sterols, as well as several vitamins^14–17^. These nutritional requirements reflect well-characterized, oxygen-dependent biosynthetic reactions. UFA synthesis involves the oxygen-dependent acyl-CoA desaturase Ole1, NAD^+^ synthesis depends on the oxygenases Bna2, Bna4, and Bna1, while synthesis of ergosterol, the main yeast sterol, even requires 12 moles of oxygen per mole.

Oxygen-dependent reactions in NAD^+^ synthesis can be bypassed by nutritional supplementation of nicotinic acid, which is a standard ingredient of synthetic media for cultivation of *S. cerevisiae*^17,18^. Ergosterol and the UFA source Tween 80 (polyethoxylated sorbitan oleate) are routinely included in media for anaerobic cultivation as ‘anaerobic growth factors’ (AGF)^15,17,19^. Under anaerobic conditions, *S. cerevisiae* imports exogenous sterols via the ABC transporters Aus1 and Pdr11^20^. Mechanisms for uptake and hydrolysis of Tween 80 by *S. cerevisiae* are unknown but, after its release, oleate is activated by the acyl-CoA synthetases Faa1 and Faa4^21,22^.

Outside the whole-genome duplicated (WGD) clade of Saccharomycotina yeasts, only few yeasts (including *Candida albicans* and *Brettanomyces bruxellensis*) are capable of anaerobic growth in synthetic media supplemented with vitamins, ergosterol and Tween 80^12,13,23,24^. However, most currently known yeast species readily ferment glucose to ethanol and carbon dioxide when exposed to oxygen-limited growth conditions^13,25,26^, indicating that they do not depend on respiration for energy conservation. The inability of the large majority of facultatively fermentative yeast species to grow under strictly anaerobic conditions is therefore commonly attributed to incompletely understood oxygen requirements for biosynthetic processes^11^. Several oxygen-requiring processes have been proposed including involvement of a respiration-coupled dihydroorotate dehydrogenase in pyrimidine biosynthesis, limitations in uptake and/or metabolism of anaerobic growth factors, and redox-cofactor balancing constraints^11,13,27^.

Quantitation, identification and elimination of oxygen requirements in non-*Saccharomyces* yeasts is hampered by the very small amounts of oxygen required for non-dissimilatory purposes. For example, preventing entry of the small amounts of oxygen required for sterol and UFA synthesis in laboratory-scale bioreactor cultures of *S. cerevisiae* requires extreme measures, such as sparging with ultra-pure nitrogen gas and use of tubing and seals that are resistant to oxygen diffusion^25,28^. This technical challenge contributes to conflicting reports on the ability of non-*Saccharomyces* yeasts to grow anaerobically, as exemplified by studies on the thermotolerant yeast *K. marxianus*^29–31^. Paradoxically, the same small oxygen requirements can represent a real challenge in large-scale bioreactors, in which oxygen availability is limited by low surface-to-volume ratios and vigorous carbon-dioxide production.

Identification of the non-dissimilatory oxygen requirements of non-conventional yeast species is required to eliminate a key bottleneck for their application in industrial anaerobic processes and, on a fundamental level, can shed light on the roles of oxygen in eukaryotic metabolism. The goal of this study was to identify and eliminate the non-dissimilatory oxygen requirements of the facultatively fermentative, thermotolerant yeast *K. marxianus*. To this end, we analyzed and compared physiological and transcriptional responses of *K. marxianus* and *S. cerevisiae* to different oxygen- and anaerobic-growth factor limitation regimes in chemostat cultures. Based on the outcome of this comparative analysis, subsequent experiments focused on characterization and engineering of sterol metabolism and yielded *K. marxianus* strains that grew anaerobically at 45 °C.

## Results

### *K. marxianus* and *S. cerevisiae* show different physiological responses to extreme oxygen limitation

To investigate oxygen requirements of *K. marxianus*, physiological responses of strain CBS6556 were studied in glucose-grown chemostat cultures operated at a dilution rate of 0.10 h^−1^ and subjected to different oxygenation and AGF limitation regimes (Fig. 1a). Physiological parameters of *K. marxianus* in these cultures were compared to those of *S. cerevisiae* CEN.PK113-7D subjected to the same cultivation regimes.

**Fig. 1.**
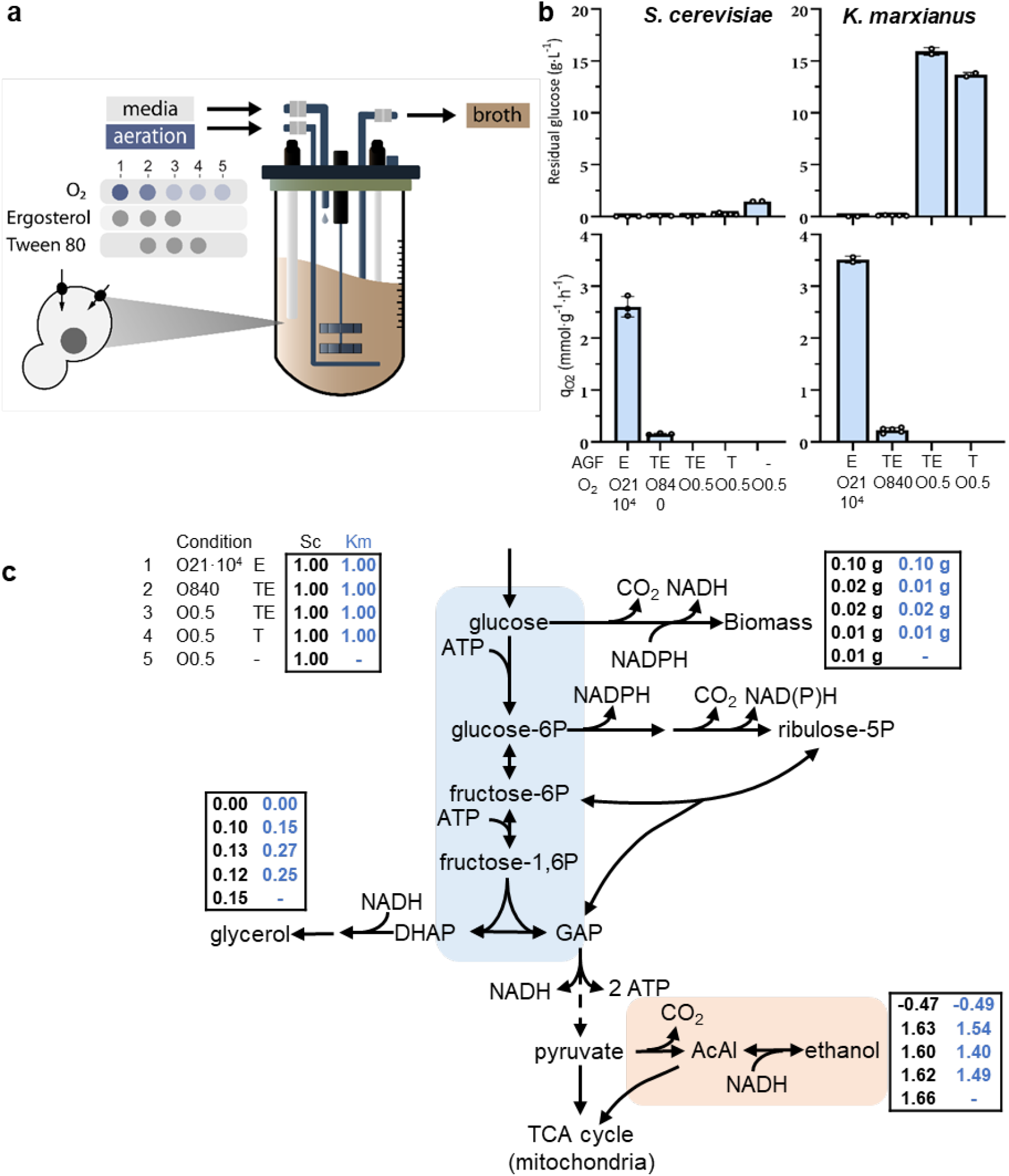
Chemostat cultivation of *S. cerevisiae* CEN.PK113-7D and *K. marxianus* CBS6556 under different aeration and anaerobic-growth-factor (AGF) supplementation regimes. The ingoing gas flow of all cultures was 500 mL·min^−1^, with oxygen partial pressures of 21·10^4^ ppm (O21·10^4^), 840 ppm (O840), or < 0.5 ppm (O0.5). The AGFs Ergosterol (E) and/or Tween 80 (T) were added to media as indicated. a, Schematic representation of experimental set-up. Data for each cultivation regime were obtained from independent replicate chemostat cultures. b, Residual glucose concentrations and biomass-specific oxygen consumption rates (q_O2_) under different aeration and AGF-supplementation regimes. Data represent mean and standard deviation of independent replicate chemostat cultures. **c**, Distribution of consumed glucose over biomass and products in chemostat cultures of *S. cerevisiae* (left column) and *K. marxianus* (right column), normalized to a glucose uptake rate of 1.00 mol·h^−1^. Numbers in boxes indicate averages of measured metabolite formation rates (mol·h^−1^) and biomass production rates (g dry weight·h^−1^) for each aeration and AGF supplementation regime.

In glucose-limited, aerobic chemostat cultures (supplied with 0.5 L air·min^−1^, corresponding to 54 mmol O_2_ h^−1^), the Crabtree-negative yeast *K. marxianus*^32^ and the Crabtree-positive yeast *S. cerevisiae*^33^ both exhibited a fully respiratory dissimilation of glucose, as evident from absence of ethanol production and a respiratory quotient (RQ) close to 1 (Table 1). Apparent biomass yields on glucose of both yeasts exceeded 0.5 g biomass (g glucose)^−1^ and were approximately 10 % higher than previously reported due to co-consumption of ethanol, which was used as solvent for the anaerobic growth factor ergosterol^32,34^.

**Table 1.**
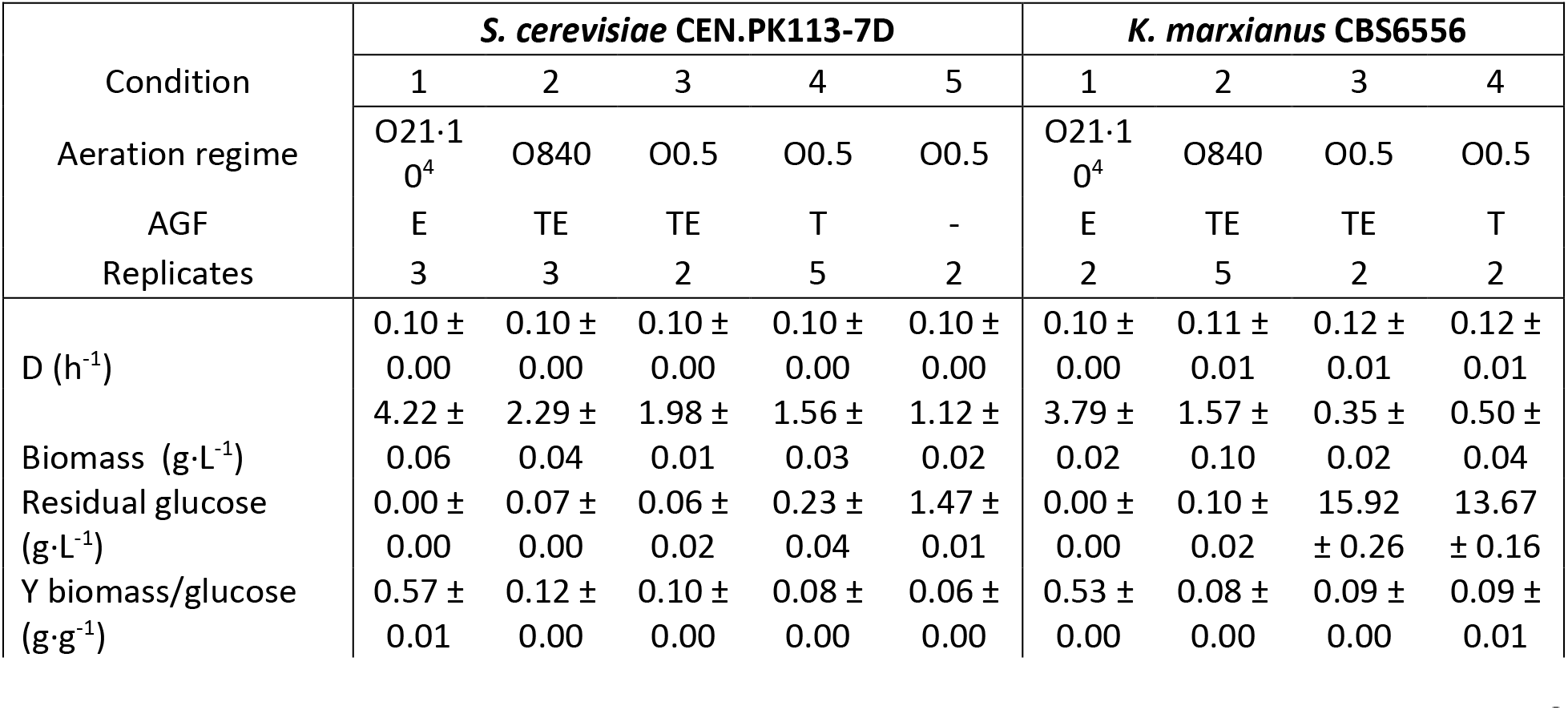
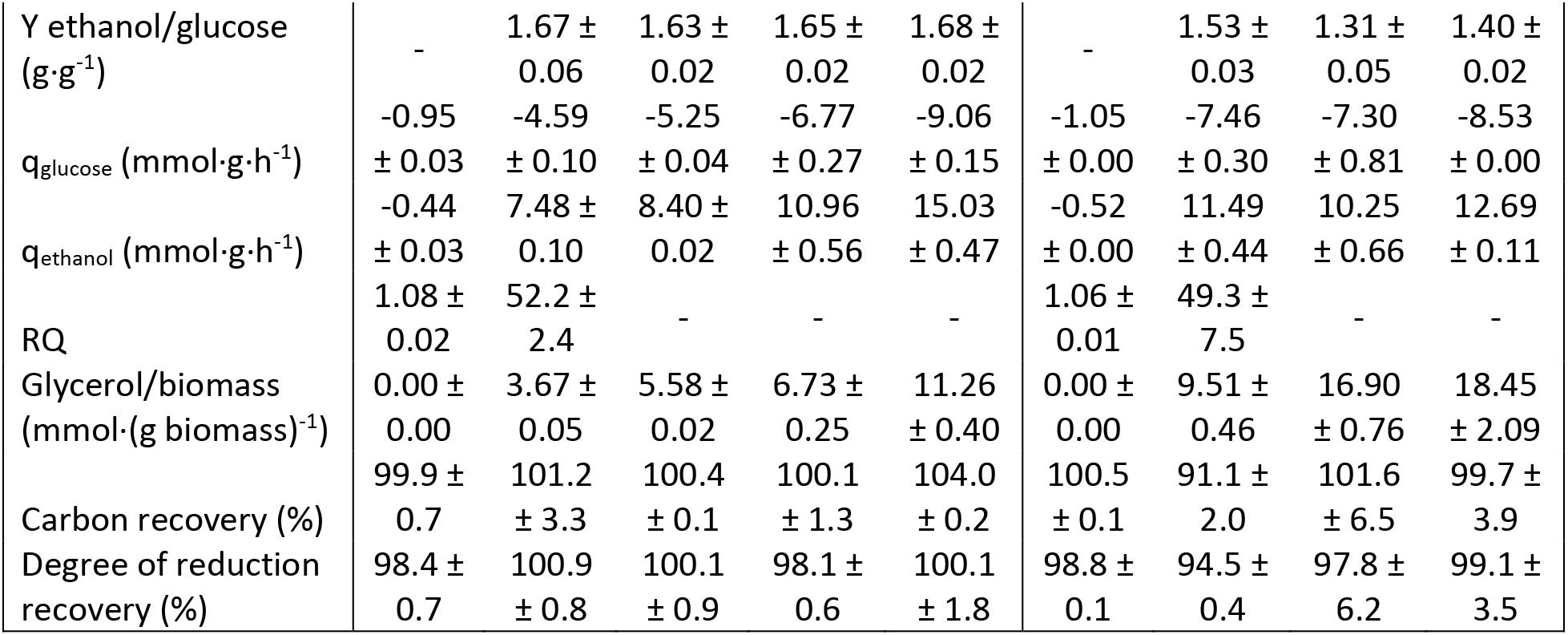
Physiology of S. cerevisiae CEN.PK113-7D and K. marxianus CBS6556 in glucose-grown chemostat cultures with different aeration and anaerobic-growth-factor (AGF) supplementation regimes. Cultures were grown at pH 6.0 on synthetic medium with urea as nitrogen source and 7.5 g·L^−1^ glucose (aerobic cultures) or 20 g·L^−1^ glucose (oxygen-limited cultures) as carbon and energy source. Data are represented as mean ± SE of data from independent chemostat cultures for each condition. The AGFs ergosterol (E) and Tween 80 (T) were added to the media as indicated. Cultures were aerated at 500 mL·min^−1^ with gas mixtures containing 21·10^4^ ppm O_2_ (O21·10^4^), 840 ppm O_2_ (O840) or < 0.5 ppm O_2_ (O0.5). Tween 80 was omitted from media used for aerobic cultivation to prevent excessive foaming. Ethanol measurements were corrected for evaporation (Supplementary Fig. 1). Positive and negative biomass-specific conversion rates (q) represent consumption and production rates, respectively.

**Table 2.** Strains used in this study. Abbreviations: Saccharomyces cerevisiae (Sc), Kluyveromyces marxianus (Km), Tetrahymena thermophila (Tt).

| Genus | Strain | Relevant genotype | Reference |
| --- | --- | --- | --- |
| *S. cerevisiae* | CEN.PK113-7D | *MATa URA3 HIS3 LEU2 TRP1 MAL2-8c SUC2* | Entian and Kötter, 2007<sup>57</sup> |
| *S. cerevisiae* | IMX585 | CEN.PK113-7D *can1Δ::cas9-natNT2* | Mans *et al.*, 2015<sup>61</sup> |
| *S. cerevisiae* | IMX1438 | IMX585 *sga1Δ::TtSTC1* | Wiersma *et al.*, 2020<sup>46</sup> |
| *S. cerevisiae* | IMK802 | IMX585 *aus1Δ* | This study |
| *S. cerevisiae* | IMK806 | IMX585 *pdr11Δ* | This study |
| *S. cerevisiae* | IMK809 | IMX585 *aus1Δ pdr11Δ* | This study |
| *K. marxianus* | CBS6556 | *URA3 HIS3 LEU2 TRP1* | CBS-KNAW* |
| *K. marxianus* | NBRC1777 | *URA3 HIS3 LEU2 TRP1* | NBRC** |
| *K. marxianus* | IMX2323 | KmPDC1p-*TtSTC1*-ScADH1t-hygB | This study |
| *K. marxianus* | IMS1111 | KmPDC1p-*TtSTC1*-ScADH1t-hygB | This study |
| *K. marxianus* | IMS1112 | KmPDC1p-*TtSTC1*-ScADH1t-hygB | This study |
| *K. marxianus* | IMS1113 | KmPDC1p-*TtSTC1*-ScADH1t-hygB | This study |
| *K. marxianus* | IMS1131 | KmPDC1p-*TtSTC1*-ScADH1t-hygB | This study |
| *K. marxianus* | IMS1132 | KmPDC1p-*TtSTC1*-ScADH1t-hygB | This study |
| *K. marxianus* | IMS1133 | KmPDC1p-*TtSTC1*-ScADH1t-hygB | This study |

**Table 3.**
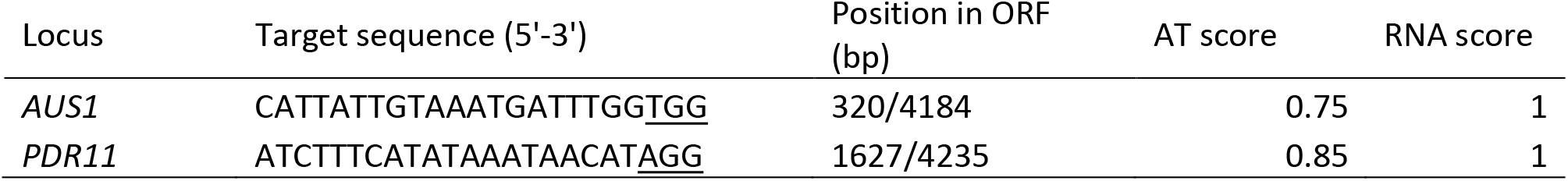
CRISPR gRNA target sequences used in this study. gRNA target sequences are shown with PAM sequences underlined. Position in ORF indicates the base pair after which the Cas9-mediated double-strand break is introduced. AT score indicates the AT content of the 20-bp target sequence and RNA score indicates the fraction of unpaired nucleotides of the 20-bp target sequence, predicted with the complete gRNA sequence using a minimum free energy prediction by the RNAfold algorithm^62^.

At a reduced oxygen-supply rate of 0.4 mmol O_2_ h^−1^, both yeasts exhibited a mixed respiro-fermentative glucose metabolism. RQ values close to 50 and biomass-specific ethanol-production rates of 11.5 ± 0.6 mmol·g·h^−1^ for *K. marxianus* and 7.5 ± 0.1 mmol·g·h^−1^ for *S. cerevisiae* (Table 1), indicated that glucose dissimilation in these cultures was predominantly fermentative. Biomass-specific rates of glycerol production which, under oxygen-limited conditions, enables re-oxidation of NADH generated in biosynthetic reactions^35^, were approximately 2.5-fold higher (*p* = *2.3·10*^−4^) in *K. marxianus* than in *S. cerevisiae*. Glycerol production showed that the reduced oxygen-supply rate constrained mitochondrial respiration. However, low residual glucose concentrations (Table 1) indicated that sufficient oxygen was provided to meet most or all of the biosynthetic oxygen requirements of *K. marxianus*.

To explore growth of *K. marxianus* under an even more stringent oxygen-limitation, we exploited previously documented challenges in achieving complete anaerobiosis in laboratory bioreactors^19,28^. Even in chemostats sparged with pure nitrogen, *S. cerevisiae* grew on synthetic medium lacking Tween 80 and ergosterol, albeit at an increased residual glucose concentration (Fig. 1, Table 1). In contrast, *K. marxianus* cultures sparged with pure N_2_ and supplemented with both AGFs consumed only 20 % of the glucose fed to the cultures. These severely oxygen-limited cultures showed a residual glucose concentration of 15.9 ± 0.3 g·L^−1^ and a low but constant biomass concentration of 0.4 ± 0.0 g·L^−1^. This pronounced response of *K. marxianus* to extreme oxygen-limitation provided an experimental context for further analyzing its unknown oxygen requirements.

*S. cerevisiae* can import exogenous sterols under severely oxygen-limited or anaerobic conditions^20^. If the latter were also true for *K. marxianus*, omission of ergosterol from the growth medium of severely oxygen-limited cultures would increase biomass-specific oxygen requirements and lead to an even lower biomass concentration. In practice however, omission of ergosterol led to a small increase of the biomass concentration and a corresponding decrease of the residual glucose concentration in severely oxygen-limited chemostat cultures (Fig. 1b, Table 1). This observation suggested that, in contrast to *S. cerevisiae*, *K. marxianus* cannot replace *de novo* oxygen-dependent sterol synthesis by uptake of exogenous sterols.

### Transcriptional responses of *K. marxianus* to oxygen limitation involve ergosterol metabolism

To further investigate the non-dissimilatory oxygen requirements of *K. marxianus*, transcriptome analyses were performed on cultures of *S. cerevisiae* and *K. marxianus* grown under the aeration and anaerobic-growth-factor supplementation regimes discussed above. The genome sequence of *K. marxianus* CBS6556 was only available as draft assembly and was not annotated^36^. Therefore, long-read genome sequencing, assembly and *de novo* genome annotation were performed, the annotation was refined by using transcriptome assemblies (**Data availability**). Comparative transcriptome analysis of *S. cerevisiae* and *K. marxianus* focused on orthologous genes with divergent expression patterns that revealed a strikingly different transcriptional response to growth limitation by oxygen and/or anaerobic-growth-factor availability (Fig. 2).

**Fig. 2.**
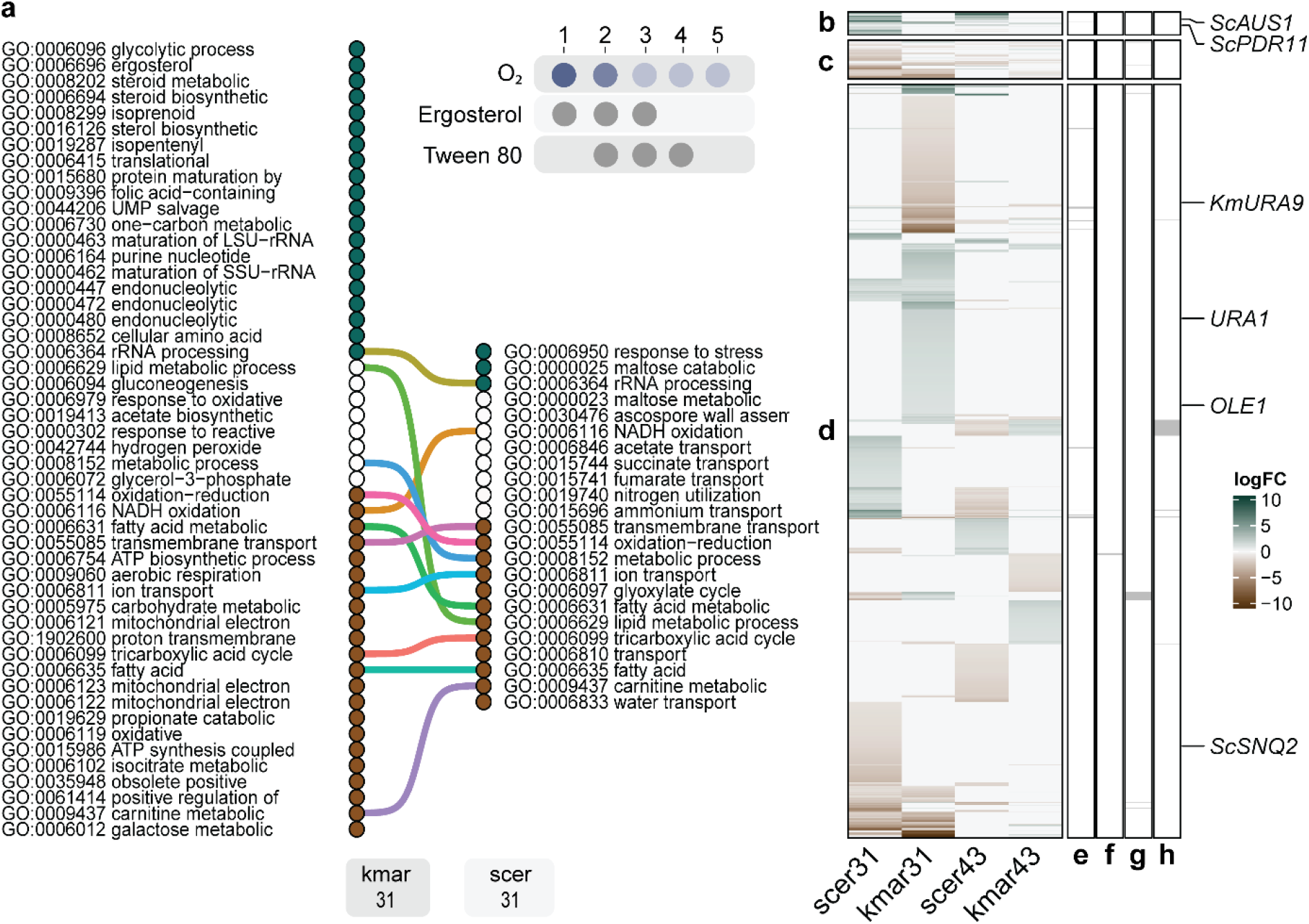
Transcriptional response of *K. marxianus* and *S. cerevisiae* to oxygen limitation and sterol, Tween 80 supplementation. Transcriptome analyses were performed for each cultivation regime (1 to 5) of *S. cerevisiae* CEN.PK113-7D (scer) and *K. marxianus* CBS6556 (kmar). Data for each regime were obtained from independent replicate chemostat cultures (Fig. 1). **a,** Comparison of GO-term gene-set enrichment analysis of biological processes in contrast 31 of *S. cerevisiae* and *K. marxianus* with short description of GO-terms (Supplementary Fig. 2-5). GO-terms were vertically ordered based on their distinct directionality calculated with Piano^40^ with GO-terms enriched solely with up-regulated genes (blue) at the top, GO-terms with mixed- or no-directionality in the middle (white) and GO-terms with solely down-regulated genes at the bottom (brown). **b**, **c**, **d**, Subsets of differentially expressed orthologous genes obtained from the gene-set analyses for both yeasts in contrasts 31 and 43, and with genes without orthologs depicted with logFC value of 0 in the respective yeast. **b,** *S. cerevisiae* genes previously shown as consistently upregulated under anaerobic conditions in four different nutrient-limitations^41^. **c**, As described for panel b but for downregulated genes. **d**, Differentially expressed genes uniquely found in this study. **e, f, g, h,** Highlighted gene-sets showing divergent expression patterns across the two yeasts. **e,** *S. cerevisiae* genes upregulated in contrast 31 but downregulated in *K. marxianus.* **f,** *S. cerevisiae* genes downregulated in contrast 31 but upregulated in *K. marxianus.* **g, h,** Similar to e and f but for contrast 43.

In *S. cerevisiae,* import of exogenous sterols by Aus1 and Pdr11 can alleviate the impact of oxygen limitation on sterol biosynthesis^20^. Consistent with this role of sterol uptake, sterol biosynthetic genes in *S. cerevisiae* were only highly upregulated in severely oxygen-limited cultures when ergosterol was omitted from the growth medium (Fig. 3b, Supplementary Fig. 6, contrast 43). Also the mevalonate pathway for synthesis of the sterol precursor squalene, which does not require oxygen, was upregulated (contrast 43), reflecting a relief of feedback regulation by ergosterol^37^. In contrast, *K. marxianus* showed a pronounced upregulation of genes involved in sterol, isoprenoid and fatty-acid metabolism (Fig. 2ab, Fig. 3, contrast 31) in severely oxygen-limited cultures supplemented with ergosterol and Tween 80. No further increase of the expression levels of sterol biosynthetic genes was observed upon omission of these anaerobic growth factors from the medium of these cultures (Supplementary Fig. 6, contrast 43). These observations suggested that *K. marxianus* may be unable to import ergosterol when sterol synthesis is compromised. Consistent with this hypothesis, co-orthology prediction with Proteinortho^38^ revealed no orthologs of the *S. cerevisiae* sterol transporters Aus1 and Pdr11 in *K. marxianus*.

**Fig. 3.**
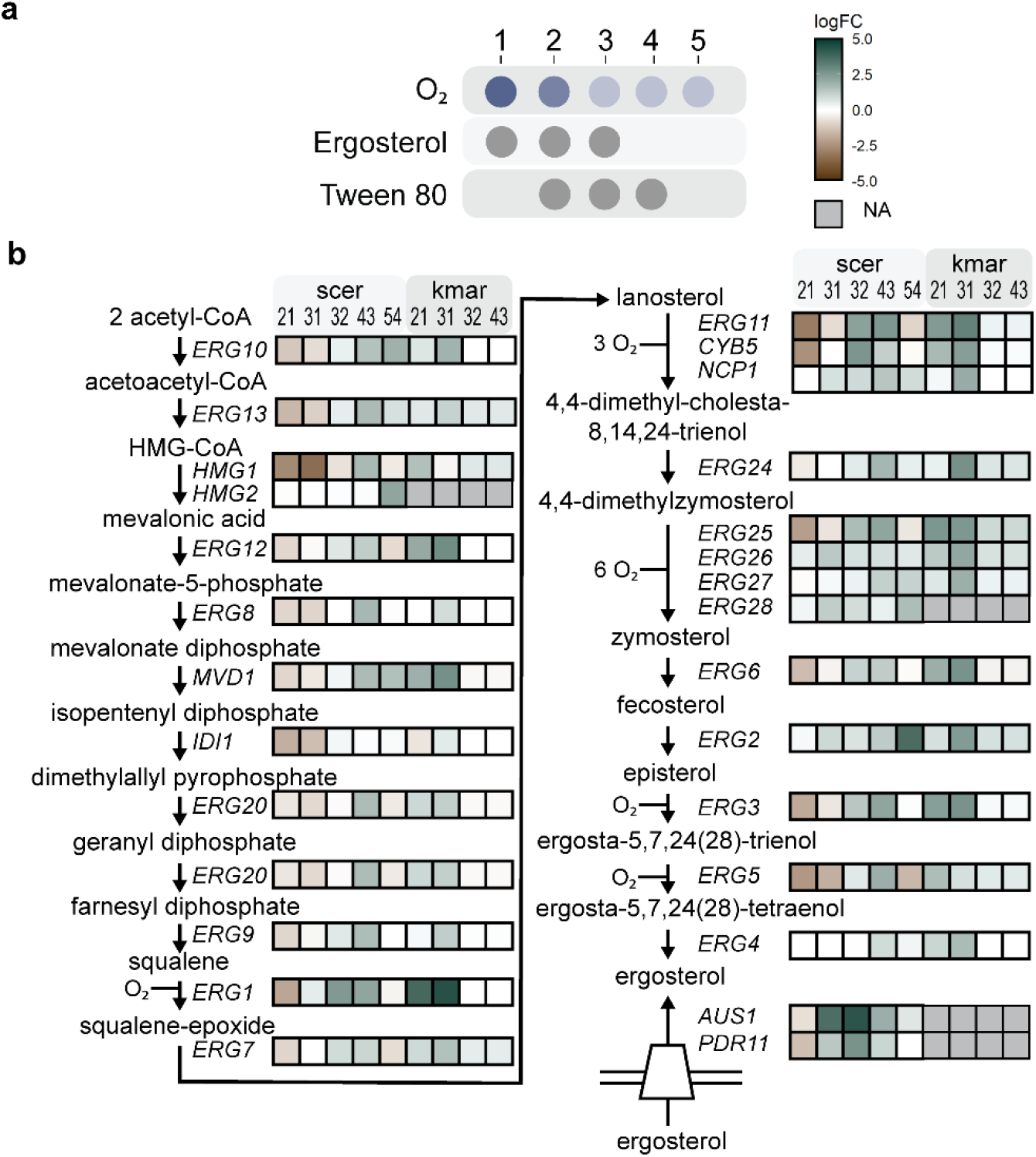
Different transcriptional regulation of ergosterol-biosynthesis in *K. marxianus* and *S. cerevisiae*. **a,** RNAseq was performed on independent replicate chemostat cultures of *S. cerevisiae* CEN.PK113-7D and *K. marxianus* CBS6556 for each aeration and anaerobic-growth-factor supplementation regime (1 to 5; Fig. 1). **b,** Transcriptional differences in the mevalonate- and ergosterol-pathway genes of *S. cerevisiae* and *K. marxianus* for contrasts 21 (O_2_ 840 TE |O 21·10^4^ E), 31 (O_2_ 0.5 TE | O 21·10^4^ E), 32 (O_2_ 0.5 TE | O_2_ 840 TE), 43 (O_2_ 0.5 T | O_2_ 0.5 TE), 54 (O_2_ 0.5 | O_2_ 0.5 T). Lumped biochemical reactions are represented by arrows. Colors indicate up- (blue) or down-regulation (brown) with color intensity indicating the log 2 fold change with color range capped to a maximum of 4. Reactions are annotated with corresponding gene, *K. marxianus* genes are indicated with the name of the *S. cerevisiae* orthologs. Ergosterol uptake by *S. cerevisiae* requires additional factors beyond the membrane transporters Aus1 and Pdr11^42^. No orthologs of the sterol-transporters or Hmg2 were identified for *K. marxianus* and low read counts for Erg3, Erg9 and Erg20 precluded differential gene expression analysis across all conditions (dark grey). Enzyme abbreviations: Erg10 acetyl-CoA acetyltransferase, Erg13 3-hydroxy-3-methylglutaryl-CoA (HMG-CoA) synthase, Hmg1/Hmg2 HMG-CoA reductase, Erg12 mevalonate kinase, Erg8 phosphomevalonate kinase, Mvd1 mevalonate pyrophosphate decarboxylase, Idi1 isopentenyl diphosphate:dimethylallyl diphosphate (IPP) isomerase, Erg20 farnesyl pyrophosphate synthetase, Erg9 farnesyl-diphosphate transferase (squalene synthase), Erg7 lanosterol synthase, Erg11 lanosterol 14α-demethylase, Cyb5 cytochrome b5 (electron donor for sterol C5-6 desaturation), Ncp1 NADP-cytochrome P450 reductase, Erg24 C-14 sterol reductase, Erg25 C-4 methyl sterol oxidase, Erg26 C-3 sterol dehydrogenase, Erg27 3-keto-sterol reductase, Erg28 endoplasmic reticulum membrane protein (may facilitate protein-protein interactions between Erg26 and Erg27, or tether these to the ER), Erg6 Δ24-sterol C-methyltransferase, Erg2 Δ24-sterol C-methyltransferase, Erg3 C-5 sterol desaturase, Erg5 C-22 sterol desaturase, Erg4 C24/28 sterol reductase, Aus1/Pdr11 plasma-membrane sterol transporter.

*K. marxianus* harbors two dihydroorotate dehydrogenases, a cytosolic fumarate-dependent enzyme (KmUra1) and a mitochondrial quinone-dependent enzyme (KmUra9). *In vivo* activity of the latter requires oxygen because the reduced quinone is reoxidized by the mitochondrial respiratory chain^39^. Consistent with these different oxygen requirements, Km*URA9* was down-regulated under severely oxygen-limited conditions, while Km*URA1* was upregulated (Fig. 2b, contrast 31). Upregulation of Km*URA1* coincided with increased production of succinate (Table 1).

### Absence of sterol import in *K. marxianus*

To test the hypothesis that *K. marxianus* lacks a functional sterol-uptake mechanism, uptake of fluorescent sterol derivative 25-NBD-cholesterol (NBDC) was measured by flow cytometry^43^. Since *S. cerevisiae* sterol transporters are not expressed in aerobic conditions^20^ and to avoid interference of sterol synthesis, NBDC uptake was analysed in anaerobic cell suspensions (Fig. 4a). Four hours after NBDC addition to cell suspensions of the reference strain *S. cerevisiae* IMX585, median single-cell fluorescence increased by 66-fold (Fig. 4bc). In contrast, the congenic sterol-transporter-deficient strain IMK809 (*aus1*Δ *pdr11*Δ) only showed a 6-fold increase of fluorescence, probably reflected detergent-resistant binding of NBDC to *S. cerevisiae* cell-wall proteins^43,44^. *K. marxianus* strains CBS6556 and NBRC1777 did not show increased fluorescence, neither after 4 h nor after 23 h of incubation with NBDC (< 2-fold, Fig. 4bc, Supplementary Fig. 7).

**Fig. 4.**
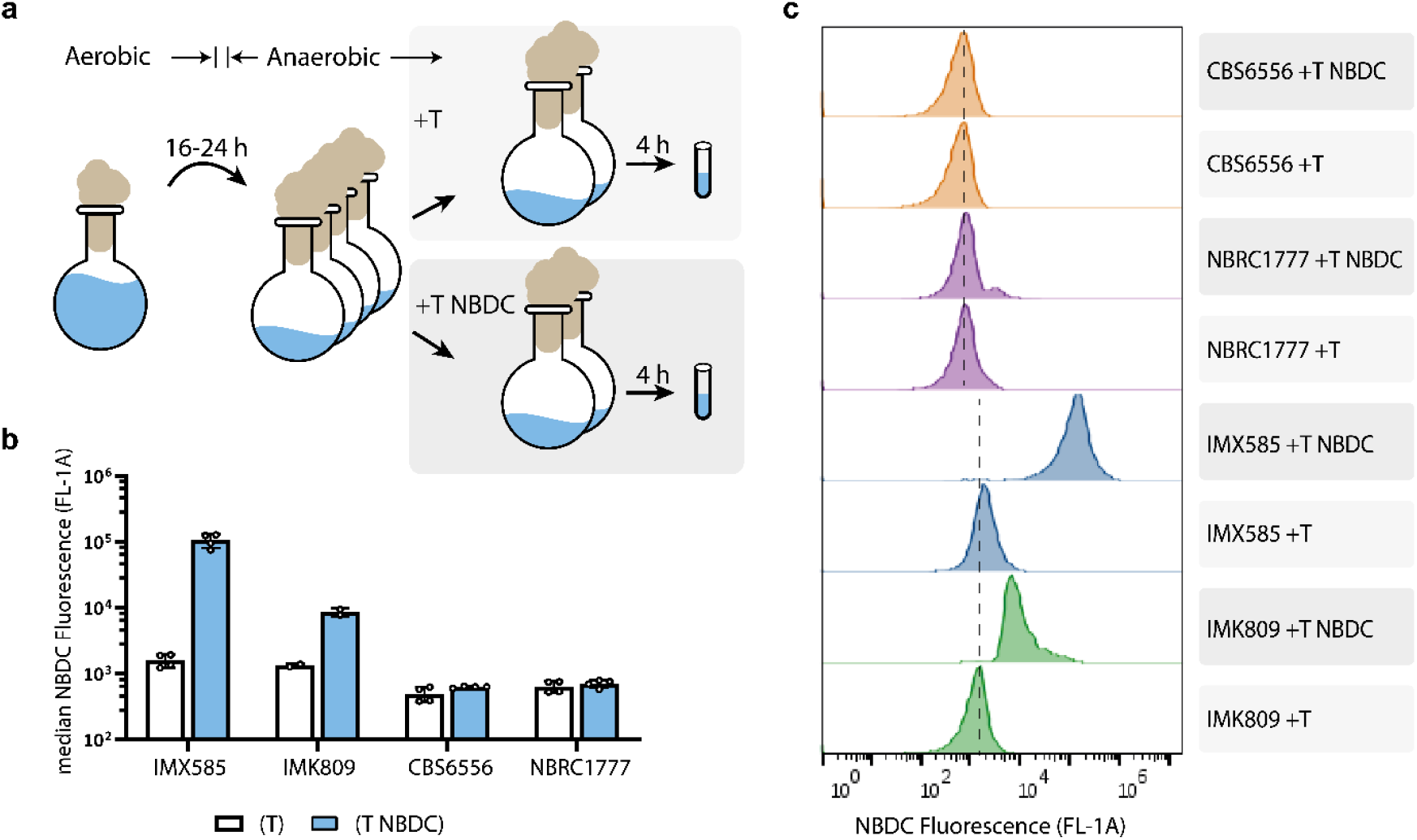
Uptake of the fluorescent sterol derivative NBDC by *S. cerevisiae* and *K. marxianus* strains. **a**, Experimental approach. *S. cerevisiae* strains IMX585 (reference) and IMK809 (*aus1*Δ *pdr11*Δ), and *K. marxianus* strains NBRC1777 and CBS6556 were each anaerobically incubated in four replicate shake-flask cultures. NBDC and Tween 80 (NBDC T) were added to two cultures, while only Tween 80 (T) was added to the other two. After 4 h incubation, cells were stained with propidium iodide (PI) and analysed by flow cytometry. PI staining was used to eliminate cells with compromised membrane integrity from analysis of NBDC fluorescence. Cultivation conditions and flow cytometry gating are described in Methods and in Supplementary Fig. 8, Supplementary Data set 1 and 2. **b**, Median and pooled standard deviation of fluorescence intensity (λ_ex_ 488 nm | λ_em_ 533/30 nm, FL1-A) of PI-negative cells with variance of biological replicates after 4 h exposure to Tween 80 (white bars) or Tween 80 and NBDC (blue bars). Variance was pooled for the strains IMX585, CBS6556 and NBRC1777 by repeating the experiment. **c**, NBDC fluorescence-intensity distribution of cells in a sample from a single culture for each strain, shown as modal-scaled density function. Dashed lines represent background fluorescence of unstained cells of *S. cerevisiae* and *K. marxianus*. Fluorescence data for 23-h incubations with NBDC are shown in Supplementary Fig. 7.

### Engineering *K. marxianus* for oxygen-independent growth

Sterol uptake by *S. cerevisiae*, which requires cell wall proteins as well as a membrane transporter, has not yet been fully resolved^42,43^. Instead of expressing a heterologous sterol-import system in *K. marxianus*, we therefore explored production of tetrahymanol, which acts as a sterol surrogate in strictly anaerobic fungi ^45^. Expression of a squalene-tetrahymanol cyclase from *Tetrahymena thermophila* (*TtSTC1*), which catalyzes the single-step oxygen-independent conversion of squalene into tetrahymanol (Fig. 5a), was recently shown to enable sterol-independent growth of *S. cerevisiae*^46^.

**Fig. 5.**
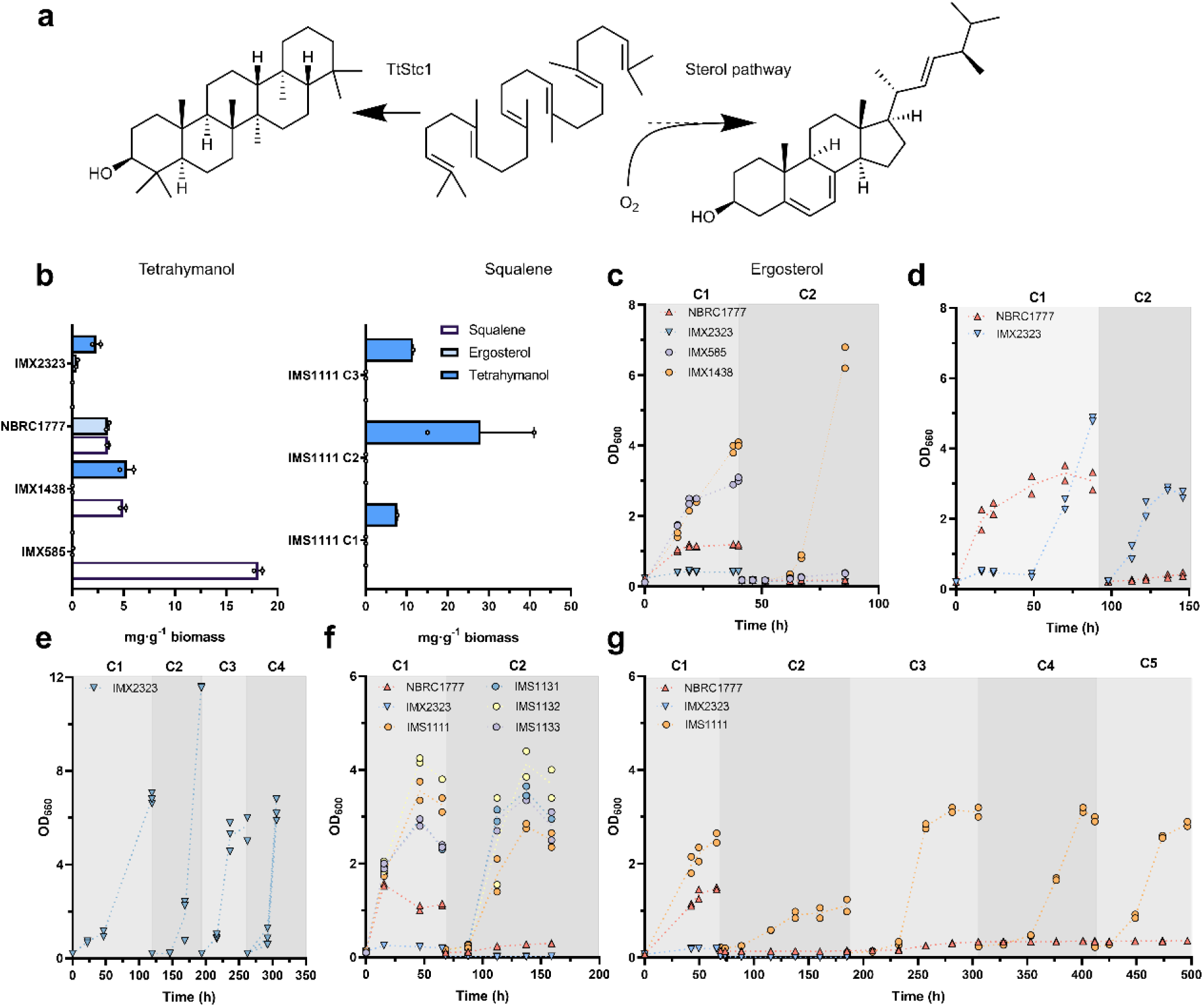
Sterol-independent anaerobic growth of *K. marxianus* strains expressing *TtSTC1*. **a**, Oxygen-dependent sterol synthesis and cyclisation of squalene to tetrahymanol by TtStc1. **b**, Squalene, ergosterol, and tetrahymanol contents with mean and standard error of the mean of (left panel) *S. cerevisiae* strains IMX585 (reference), IMX1438 (*sga1*Δ::*TtSTC1*), and *K. marxianus* strains NBRC1777 (reference), IMX2323 (*TtSTC1*). Lipid composition of single-cell isolate IMS1111 (*TtSTC1*) (right panel) over 3 serial transfers (C1-C3). Data from replicate cultures grown in strictly anaerobic (**c**, **f**, **g**) or severely oxygen-limited shake-flask cultures (**d, e**). Aerobic grown pre-cultures were used to inoculate the first anaerobic culture on SMG-urea and Tween 80, when the optical density started to stabilize the cultures were transferred to new media. Data depicted are of each replicate culture (points) and the mean (dotted line) from independent biological duplicate cultures, serial transfers cultures are represented with C1-C5. Strains NBRC1777 (wild-type, upward red triangles), IMX2323 (*TtSTC1*, cyan downward triangle), and the single-cell isolates IMS1111 (*TtSTC1*, orange circles), IMS1131 (*TtSTC1*, blue circles), IMS1132 (*TtSTC1*, yellow circles), IMS1133 (*TtSTC1*, purple circles). *S. cerevisiae* IMX585 (reference, purple circle) and IMX1438 (*TtSTC1*, orange circles). **c**, Extended data with double inoculum size is available in Supplementary Fig. 10. **d**, Extended data is available in Supplementary Fig. 9a.

*TtSTC1* was expressed in *K. marxianus* NBRC1777, which is more genetically amenable than strain CBS6556^47^. After 40 h of anaerobic incubation, the resulting strain contained 2.4 ± 0.4 mg·(g biomass)^−1^ tetrahymanol, 0.4 ± 0.1 mg·g^−1^ ergosterol and no detectable squalene, while strain NBRC1777 contained 3.5 ± 0.1 mg·g^−1^ squalene and 3.4 ± 0.2 mg·g^−1^ ergosterol (Fig. 5b). In strictly anaerobic cultures on sterol-free medium, strain NBRC1777 grew immediately after inoculation but not after transfer to a second anaerobic culture (Fig. 5c), consistent with ‘carry-over’ of ergosterol from the aerobic preculture^19^. The tetrahymanol-producing strain did not grow under these conditions (Fig. 5c) but showed sustained growth under severely oxygen-limited conditions that did not support growth of strain NBRC1777 (Fig. 5de). Single-cell isolates derived from these oxygen-limited cultures (IMS1111, IMS1131, IMS1132, IMS1133) showed instantaneous as well as sustained growth under strictly anaerobic conditions (Figure 5f and 5g). Tetrahymanol contents in the first, second and third cycle of anaerobic cultivation of isolate IMS1111 were 7.6 ± 0.0 mg·g^−1^, 28.0 ± 13.0 mg·g^−1^ and 11.5 ± 0.1 mg·g^−1^, respectively (Fig. 5b), while no ergosterol was detected.

To identify whether adaptation of the tetrahymanol-producing strain IMX2323 to anaerobic growth involved genetic changes, its genome and those of the four adapted isolates were sequenced (Supplementary Table 1). No copy number variations were detected in any of the four adapted isolates. Only strain IMS1111 showed two non-conservative mutations in coding regions: a single-nucleotide insertion in a transposon-borne gene and a stop codon at position 350 (of 496 bp) in *KmCLN3*, which encodes for a G1 cyclin^48^. The apparent absence of mutations in the three other, independently adapted strains indicated that their ability to grow anaerobically reflected a non-genetic adaptation.

### Test of anaerobic thermotolerance and selection for fast growing anaerobes

One of the attractive phenotypes of *K. marxianus* for industrial application is its high thermotolerance with reported maximum growth temperatures of 46-52 °C^49,50^. To test if anaerobically growing tetrahymanol-producing strains retained thermotolerance, strain IMS1111 was grown in anaerobic sequential-batch-reactor (SBR) cultures (Fig. 6) in which, after an initial growth cycle at 30 °C, the growth temperature was shifted to 42 °C. Specific growth at 42 °C progressively accelerated from 0.06 h^−1^ to 0.13 h^−1^ over 17 SBR cycles (corresponding to ca. 290 generations; Fig. 6b). A subsequent temperature increase to 45 °C led to a strong decrease of the specific growth rate which, after approximately 1000 generations of selective growth, stabilized at approximately 0.08 h^−1^. Whole-population genome sequencing of the evolved populations revealed no common mutations or chromosomal copy number variations (Supplementary Table 1). These data show that *TtSTC1*-expressing *K. marxianus* can grow anaerobically at temperatures up to at least 45 °C.

**Fig. 6.**
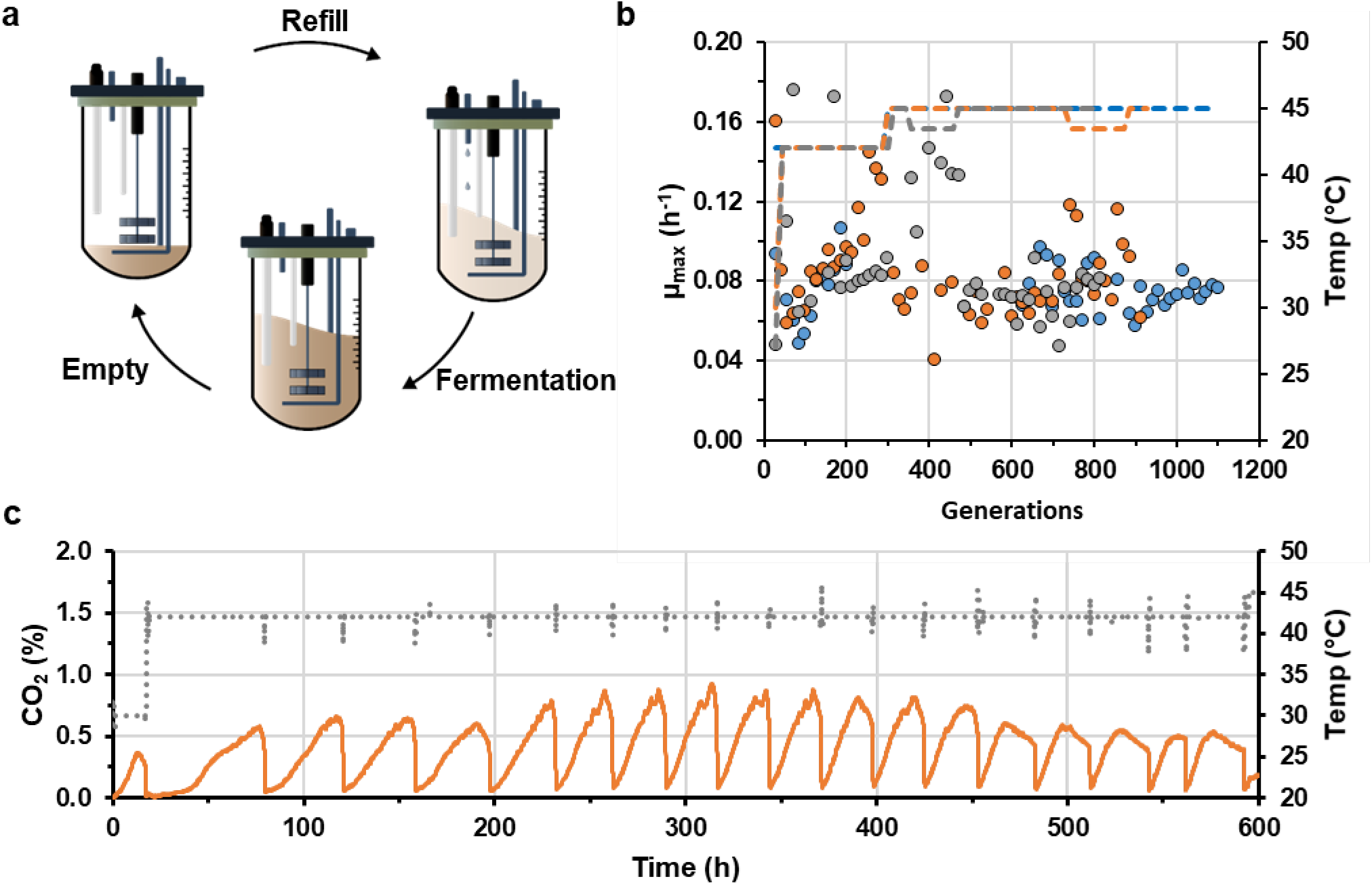
Thermotolerance and anaerobic growth of tetrahymanol-producing *K. marxianus* strain. The strain IMS1111 was grown in triplicate sequential batch bioreactor cultivations in synthetic media supplemented with 20 g·L^−1^ glucose and 420 mg·L^−1^ Tween 80 at pH 5.0. **a**, Experimental design of sequential batch fermentation with cycles at step-wise increasing temperatures to select for faster growing mutants, each cycle consisted of three phases; (i) (re)filling of the bioreactor with fresh media up to 100 mL and adjustment of temperature to a new set-point, (ii) anaerobic batch fermentation at a fixed culture temperature with continuous N_2_ sparging for monitoring of CO_2_ in the culture off-gas, and (iii) fast broth withdrawal leaving 7 mL (14.3 fold dilution) to inoculate the next batch. **b**, Maximum specific estimated growth rate (circles) of each batch cycle for the three independent bioreactor cultivations (M3R blue, M5R orange, M6L grey) with the estimated number of generations. The growth rate was calculated from the CO_2_ production as measured in the off-gas and should be interpreted as an estimate and in some cases could not be calculated. The culture temperature profile (dotted line) for each independent bioreactor cultivation (blue, grey, orange) consisted of a step-wise increment of the temperature at the onset of the fermentation phase in each batch cycle. **c**, Representative section of CO_2_ off-gas profiles of the individual bioreactor (M5R) cultivation over time with CO_2_ fraction (orange line) and culture temperature (grey dotted line), data of the entire experiment is available in Supplementary Fig. 11 (Data availability).

## Discussion

Industrial production of ethanol from carbohydrates relies on *S. cerevisiae*, due to its capacity for efficient, fast alcoholic fermentation and growth under strictly anaerobic process conditions. Many facultatively fermentative yeast species outside the Saccharomycotina WGD-clade also rapidly ferment sugars to ethanol under oxygen-limited conditions^26^, but cannot grow and ferment in the complete absence of oxygen^11,13,25^. Identifying and eliminating oxygen requirements of these yeasts is essential to unlock their industrially relevant traits for application. Here, this challenge was addressed for the thermotolerant yeast *K. marxianus*, using a systematic approach based on chemostat-based quantitative physiology, genome and transcriptome analysis, sterol-uptake assays and genetic modification. *S. cerevisiae*, which was used as a reference in this study, shows strongly different genome-wide expression profiles under aerobic and anaerobic or oxygen-limited conditions^51^. Although only a small fraction of these differences were conserved in *K. marxianus* (Fig. 2), we were able to identify absence of a functional sterol import system as the critical cause for its inability to grow anaerobically. Enabling synthesis of the sterol surrogate tetrahymanol yielded strains that grew anaerobically at temperatures above the permissive temperature range of *S. cerevisiae*.

A short adaptation phase of tetrahymanol-producing *K. marxianus* strains under oxygen-limited conditions reproducibly enabled strictly anaerobic growth. Although this ability was retained after aerobic isolation of single-cell lines, we were unable to attribute this adaptation to mutations. In contrast to wild-type *K. marxianus*, a non-adapted tetrahymanol-producing strain did not show ‘carry-over growth’ after transfer from aerobic to strictly anaerobic conditions and adapted cultures showed reduced squalene contents (Fig. 5). These observations suggest that interactions between tetrahymanol, ergosterol and/or squalene influence the onset of anaerobic growth and that oxygen-limited growth results in a stable balance between these lipids that is permissive for anaerobic growth.

Comparative genomic studies in Saccharomycotina yeasts have previously led to the hypothesis that sterol transporters are absent from pre-WGD yeast species^11,52^. While our observations on *K. marxianus* reinforce this hypothesis, which was hitherto not experimentally tested, they do not exclude involvement of additional oxygen-requiring reactions in other non-*Saccharomyces* yeasts. For example, pyrimidine biosynthesis is often cited as a key oxygen-requiring process in non-*Saccharomyces* yeasts, due to involvement of a respiratory-chain-linked dihydroorotate dehydrogenase (DHOD)^53,54^. *K. marxianus*, is among a small number of yeast species that, in addition to this respiration dependent enzyme (KmUra9), also harbors a fumarate-dependent DHOD (KmUra1)^55^. In *K. marxianus* the activation of this oxygen-independent KmUra1 is a crucial adaptation for anaerobic pyrimidine biosynthesis. The experimental approach followed in the present study should be applicable to resolve the role of pyrimidine biosynthesis and other oxygen-requiring reactions in additional yeast species.

Enabling *K. marxianus* to grow anaerobically represents an important step towards application of this thermotolerant yeast in large-scale anaerobic bioprocesses. However, specific growth rates and biomass yields of tetrahymanol-expressing *K. marxianus* in anaerobic cultures were lower than those of wild-type *S. cerevisiae* strains. A similar phenotype of tetrahymanol-producing *S. cerevisiae* was proposed to reflect an increased membrane permeability^46^. Additional membrane engineering or expression of a functional sterol transport system is therefore required for further development of robust, anaerobically growing industrial strains of *K. marxianus*^56^.

## online Methods

### Yeast strains, maintenance and shake-flask cultivation

*Saccharomyces cerevisiae* CEN.PK113-7D^57,58^ (*MATa MAL2-8c SUC2*) was obtained from Dr. Peter Kötter, J.W. Goethe University, Frankfurt. *Kluyveromyces marxianus* strains CBS 6556 (ATCC 26548; NCYC 2597; NRRL Y-7571) and NBRC 1777 (IFO 1777) were obtained from the Westerdijk Fungal Biodiversity Institute (Utrecht, The Netherlands) and the Biological Resource Center, NITE (NBRC) (Chiba, Japan), respectively. Stock cultures of *S. cerevisiae* were grown at 30 °C in an orbital shaker set at 200 rpm, in 500 mL shake flasks containing 100 mL YPD (10 g·L^−1^ Bacto yeast extract, 20 g·L^−1^ Bacto peptone, 20 g·L^−1^ glucose). For cultures of *K. marxianus*, the glucose concentration was reduced to 7.5 g·L^−1^. After addition of glycerol to early stationary-phase cultures, to a concentration of 30 % (v/v), 2 mL aliquots were stored at −80 °C. Shake-flask precultures for bioreactor experiments were grown in 100 mL synthetic medium (SM) with glucose as carbon source and urea as nitrogen source (SMG-urea)^17,59^. For anaerobic cultivation, synthetic medium was supplemented with ergosterol (10 mg·L^−1^) and Tween 80 (420 mg·L^−1^) as described previously^14,17,19^.

### Expression cassette and plasmid construction

Plasmids used in this study are described in (Table 4). To construct plasmids pUDE659 (gRNA*_AUS1_*) and pUDE663 (gRNA*_PDR11_*), the pROS11 plasmid-backbone was PCR amplified using Phusion HF polymerase (Thermo Scientific, Waltham, MA) with the double-binding primer 6005. PCR amplifications were performed with desalted or PAGE-purified oligonucleotide primers (Sigma-Aldrich, St Louis, MO) according to manufacturer’s instructions. To introduce the gRNA-encoding nucleotide sequences into gRNA-expression plasmids, a 2μm fragment was first amplified with primers 11228 and 11232 containing the specific sequence as primer overhang using pROS11 as template. PCR products were purified with genElutePCR Clean-Up Kit (Sigma-Aldrich) or Gel DNA Recovery Kit (Zymo Research, Irvine, CA). The two DNA fragments were then assembled by Gibson Assembly (New England Biolabs, Ipswich, MA) according to the manufacturer’s instructions. Gibson assembly reaction volumes were downscaled to 10 µL and 0.01 pmol·µL^−1^ DNA fragments at 1:1 molar ratio for 1 h at 50 °C. Chemically competent *E. coli* XL1-Blue was transformed with the Gibson assembly mix via a 5 min incubation on ice followed by a 40 s heat shock at 42 °C and 1 h recovery in non-selective LB medium. Transformants were selected on LB agar containing the appropriate antibiotic. Golden Gate assembly with the yeast tool kit^60^ was performed in 20 µL reaction mixtures containing 0.75 µL BsaI HF V2 (NEB, #R3733), 2 µL DNA ligase buffer with ATP (New England Biolabs), 0.5 µL T7-ligase (NEB) with 20 fmol DNA donor fragments and MilliQ water. Before ligation at 16 °C was initiated by addition of T7 DNA ligase, an initial BsaI digestion (30 min at 37 °C) was performed. Then 30 cycles of digestion and ligation at 37 °C and 16 °C, respectively, were performed, with 5 min incubation times for each reaction. Thermocycling was terminated with a 5 min final digestion step at 60 °C.

**Table 4.**
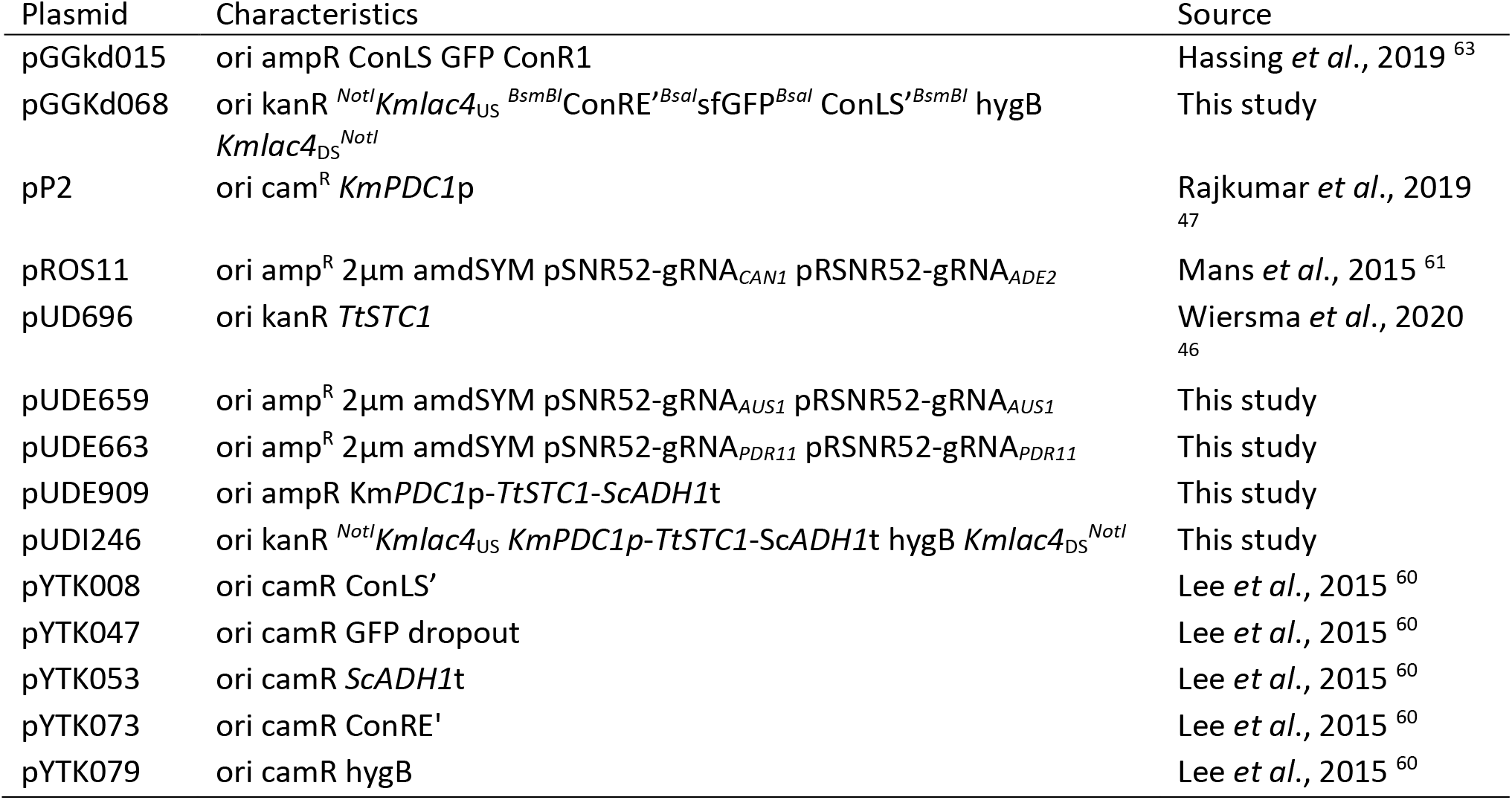
Plasmids used in this study. Restriction enzyme recognition sites are indicated in superscript. US/DS represent upstream and downstream homologous recombination sequences used for genomic integration into the *K. marxianus lac4* locus. Abbreviations: *Saccharomyces cerevisiae* (Sc), *Kluyveromyces marxianus* (Km), *Tetrahymena thermophila* (Tt).

**Table 5.**
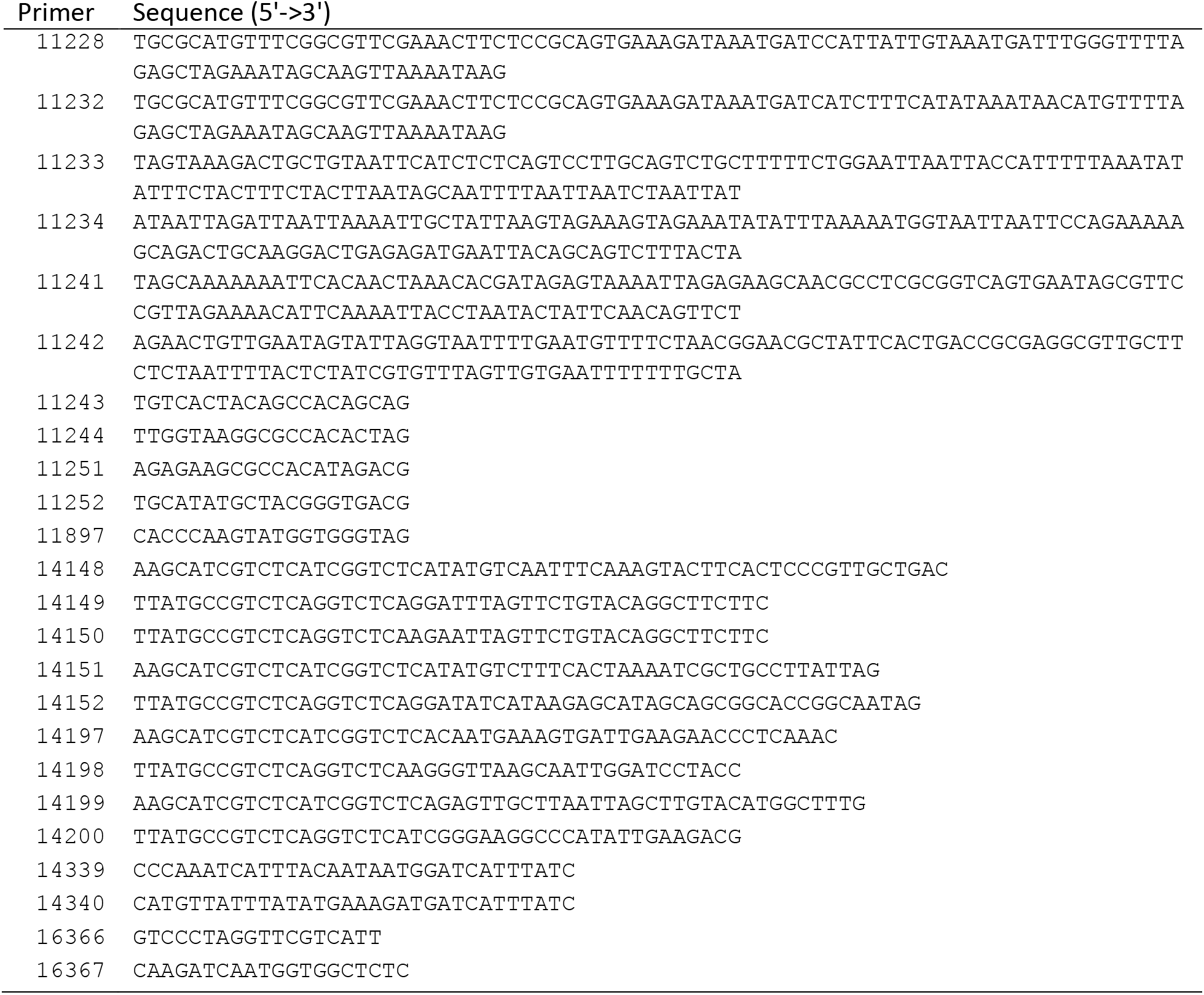
Oligonucleotide primers used in this study.

To construct a *TtSTC1* expression vector, the coding sequence of *TtSTC1* (pUD696) was PCR amplified with primer pair 16096/16097 and Golden gate assembled with the donor plasmids pGGkd015 (ori ampR), pP2 (Km*PDC1*p), pYTK053 (Sc*ADH1*t) resulting in pUDE909 (ori ampR Km*PDC1*p-*TtSTC1*-Sc*ADH1*t). For integration of *TtSTC1* cassette into the *lac4* locus both upstream and downstream flanks (877/878 bps) of the *lac4* locus were PCR amplified with the primer pairs 14197/14198 and 14199/14200, respectively. An empty integration vector, pGGKd068, was constructed by *BsaI* golden gate cloning of pYTK047 (GFP-dropout), pYTK079 (hygB), pYTK090 (kanR), pYTK073 (ConRE’), pYTK008 (ConLS’) together with the two *lac4* homologous nucleotide sequences. Plasmid assembly was verified by PCR amplification with primers 15210, 9335, 16274 and 16275 and by digestion with *BsmBI* (New England Biolabs, #R0580). The integration vector pUDI246 with the *TtSTC1* expression cassette was constructed by Gibson assembly of the PCR amplified pGGKd068 and pUDE909 with primer pairs 16274/16275 and 16272/16273, thereby adding 20 bp overlaps for assembly. For this step, the incubation time of the Gibson assembly was increased to 90 min. Plasmid assembly was verified by diagnostic PCR amplification using DreamTaq polymerase (Thermo Scientific) with primers 5941, 8442, 15216 and subsequent Illumina short-read sequencing.

### Strain construction

The lithium-acetate/polyethylene-glycol method was used for yeast transformation^64^. Homologous repair (HR) DNA fragments for markerless CRISPR-Cas9-mediated gene deletions in *S. cerevisiae* were constructed by annealing two 120 bp primers, using primer pairs 11241/11242 and 11233/11234 for deletion of *PDR11* and *AUS1*, respectively. After transformation of *S. cerevisiae* IMX585 with gRNA plasmids pUDE659 and pUDE663 and double-stranded repair fragments, transformants were selected on synthetic medium with acetamide as sole nitrogen source^65^. Deletion of *AUS1* and *PDR11* was confirmed by PCR amplification with primer pairs 11243/11244 and 11251/11252, respectively. Loss of gRNA plasmids was induced by cultivation of single-colony isolates on YPD, after which plasmid loss was assessed by absence of growth of single-cell isolates on synthetic medium with acetamide as nitrogen source. An *aus1Δ pdr11Δ* double-deletion strain was similarly constructed by chemical transformation of *S. cerevisiae* IMK802 with pUDE663 and repair DNA. To integrate a *TtSTC1* expression cassette into the *K. marxianus lac4* locus, *K. marxianus* NBRC1777 was transformed with 2 μg DNA *NotI*-digested pUDI246. After centrifugation, cells were resuspended in YPD and incubated at 30 °C for 3 h. Cells were then again centrifuged, resuspended in demineralized water and plated on 200 µg·L^−1^ hygromycin B (InvivoGen, Toulouse, France) containing agar with 40 µg·L^−1^ X-gal, 5-bromo-4-chloro-3-indolyl-β-D-galactopyranoside (Fermentas, Waltham, MA). Colonies that could not convert X-gal were analyzed for correct genomic integration of the *TtSTC1* by diagnostic PCR with primers 16366, 16367 and 11897. Genomic integration of *TtSTC1* into the chromosome outside the *lac4* locus was confirmed by short-read Illumina sequencing.

### Chemostat cultivation

Chemostat cultures were grown at 30 °C in 2 L bioreactors (Applikon, Delft, the Netherlands) with a stirrer speed of 800 rpm. The dilution rate was set at 0.10 h^−1^ and a constant working volume of 1.2 L was maintained by connecting the effluent pump to a level sensor. Cultures were grown on synthetic medium with vitamins^17^. Concentrated glucose solutions were autoclaved separately at 110 °C for 20 min and added at the concentrations indicated, along with sterile antifoam pluronic 6100 PE (BASF, Ludwigshafen, Germany; final concentration 0.2 g·L^−1^). Before autoclaving, bioreactors were tested for gas leakage by submerging them in water while applying a 0.3 bar overpressure.

Anaerobic conditions of bioreactor cultivations were maintained by continuous reactor headspace aeration with pure nitrogen gas (≤ 0.5 ppm O_2_, HiQ Nitrogen 6.0, Linde AG, Schiedam, the Netherlands) at a flowrate of 500 mL N_2_ min^−1^ (2.4 vvm). Gas pressure of 1.2 bar of the reactor headspace was set with a reduction valve (Tescom Europe, Hannover, Germany) and remained constant during cultivation. To prevent oxygen diffusion into the cultivation the bioreactor was equipped with Fluran tubing (14 Barrer O_2_, F-5500-A, Saint-Gobain, Courbevoie, France), Viton O-rings (Eriks, Alkmaar, the Netherlands), and no pH probes were mounted. The medium reservoir was deoxygenated by sparge aeration with nitrogen gas (≤ 1 ppm O_2_, HiQ Nitrogen 5.0, Linde AG).

For aerobic cultivation the reactor was sparged continuously with dried air at a flowrate of 500 mL air min^−1^ (2.4 vvm). Dissolved oxygen levels were analyzed by Clark electrodes (AppliSens, Applikon) and remained above 40% during the cultivation. For micro-aerobic cultivations nitrogen (≤ 1 ppm O_2_, HiQ Nitrogen 5.0, Linde AG) and air were mixed continuously by controlling the fractions of mass flow rate of the dry gas to a total flow of 500 mL min^−1^ per bioreactor. The mixed gas was distributed to each bioreactor and analyzed separately in real-time. Continuous cultures were assumed to be in steady state when after at least 5 volumes changes, culture dry weight and the specific carbon dioxide production rates changed by less than 10%.

Cell density was routinely measured at a wavelength of 660 nm with spectrophotometer Jenway 7200 (Cole Palmer, Staffordshire, UK). Cell dry weight of the cultures were determined by filtering exactly 10 mL of culture broth over pre-dried and weighed membrane filters (0.45 µm, Thermo Fisher Scientific), which were subsequently washed with demineralized water, dried in a microwave oven (20 min, 350 W) and weighed again^66^.

### Metabolite analysis

For determination of substrate and extracellular metabolite concentrations, culture supernatants were obtained by centrifugation of culture samples (5 min at 13000 rpm) and analyzed by high-performance liquid chromatography (HPLC) on a Waters Alliance 2690 HPLC (Waters, MA, USA) equipped with a Bio-Rad HPX-87H ion exchange column (BioRad, Veenendaal, the Netherlands) operated at 60 °C with a mobile phase of 5 mM H_2_SO_4_ at a flowrate of 0.6 mL·min^−1^. Compounds were detected by means of a dual-wavelength absorbance detector (Waters 2487) and a refractive index detector (Waters 2410) and compared to reference compounds (Sigma-Aldrich). Residual glucose concentrations in continuous cultivations were determined by HPLC analysis from rapid quenched culture samples with cold steel beads^67^.

### Gas analysis

The off-gas from bioreactor cultures was cooled with a condenser (2 °C) and dried with PermaPure Dryer (Inacom Instruments, Veenendaal, the Netherlands) prior to analysis of the carbon dioxide and oxygen fraction with a Rosemount NGA 2000 Analyser (Baar, Switzerland). The Rosemount gas analyzer was calibrated with defined mixtures of 1.98 % O_2_, 3.01 % CO_2_ and high quality nitrogen gas N6 (Linde AG).

### Ethanol evaporation rate

To correct for ethanol evaporation in the continuous bioreactor cultivations the ethanol evaporation rate was determined in the same experimental bioreactor set-up without the yeast. To SM glucose media with urea 400 mM of ethanol was added after which the decrease in the ethanol concentration was measured over time by periodic measurements and quantification by HPLC analysis over the course of at least 140 hours. To reflect the media composition used for the different oxygen regimes and anaerobic growth factor supplementation, the ethanol evaporation was measured for bioreactor sparge aeration with Tween 80, bioreactor head-space aeration both with and without Tween 80. The ethanol evaporation rate was measured for each condition in triplicate.

### Lipid extractions & GC analysis

For analysis of triterpene and triterpenoid cell contents biomass was harvested, washed once with demineralized water and stored as pellet at −80 °C before freeze-drying the pellets using an Alpha 1-4 LD Plus (Martin Christ, Osterode am Harz, Germany) at −60 °C and 0.05 mbar. Freeze-dried biomass was saponificated with 2.0 M NaOH (Bio-Ultra, Sigma-Aldrich) in methylation glass tubes (PYREX^TM^ Boroslicate glass, Thermo Fisher Scientific) at 70 °C. As internal standard 5α-cholestane (Sigma-Aldrich) was added to the saponified biomass suspension. Subsequently tert-butyl-methyl-ether (tBME, Sigma-Aldrich) was added for organic phase extraction. Samples were extracted twice using tBME and dried with sodium-sulfate (Merck, Darmstadt, Germany) to remove remaining traces of water. The organic phase was either concentrated by evaporation with N_2_ gas aeration or transferred directly to an injection vial (VWR International, Amsterdam, the Netherlands). The contents were measured by GC-FID using Agilent 7890A Gas Chromatograph (Agilent Technologies, Santa Clara, CA) equipped with an Agilent CP9013 column (Agilent). The oven was programmed to start at 80 °C for 1 min, ramp first to 280 °C with 60 °C·min^−1^ and secondly to 320 °C with a rate of 10 °C·min^−1^ with a final temperature hold of 15 min. Spectra were compared to separate calibration lines of squalene, ergosterol, α-cholestane, cholesterol and tetrahymanol as described previously^46^.

### Sterol uptake assay

Sterol uptake was monitored by the uptake of fluorescently labelled 25-NBD-cholesterol (Avanti Polar Lipids, Alabaster, AL). A stock solution of 25-NBD-cholesterol (NBDC) was prepared in ethanol under an argon atmosphere and stored at −20 °C. Shake flasks with 10 mL SM glucose media were inoculated with yeast strains from a cryo-stock and cultivated aerobically at 200 rpm at 30 °C overnight. The yeast cultures were subsequently diluted to an OD_660_ of 0.2 in 400 mL SM glucose media in 500 mL shake flasks to gradually reduce the availability of oxygen and incubated overnight. Yeast cultures were transferred to fresh SM media with 40 g·L^−1^ glucose and incubated under anaerobic conditions at 30 °C at 200 rpm. After 22 hours of anaerobic incubation 4 µg·L^−1^ NBD-cholesterol with 420 mg·L^−1^ Tween 80 were pulsed to the cultures. Samples were taken and washed with PBS 5 mL·L^−1^ Tergitol NP-40 pH 7.0 (Sigma-Aldrich) twice before resuspension in PBS and subsequent analysis. Propidium Iodide (PI) (Invitrogen) was added to the sample (20 µM) and stained according to the manufacturer’s instructions^68^. PI intercalates with DNA in cells with a compromised cell membrane, which results in red fluorescence. Samples both unstained and stained with PI were analyzed with Accuri C6 flow cytometer (BD Biosciences, Franklin Lakes, NJ) with a 488 nm laser and fluorescence was measured with emission filter of 533/30 nm (FL1) for NBD-cholesterol and > 670 nm (FL3) for PI. Cell gating and median fluorescence of cells were determined using FlowJo (v10, BD Bioscience). Cells were gated based on forward side scatter (FSC) and side-scatter (SSC) to exclude potential artifacts or clumping cells. Within this gated population PI positive and negatively stained cells were differentiated based on the cell fluorescence across a FL3 FL1 dimension. Flow cytometric gates were drafted for each yeast species and used for all samples. The gating strategy is given in Supplementary Fig. 8. Fluorescence of a strain was determined by a sample of cells from independent shake-flask cultures and compared to cells from identical unstained cultures of cells with the exact same chronological age. The staining experiment of the strains IMX585, CBS6556 and NBRC1777 samples was repeated twice for reproducibility, the mean and pooled variance was subsequently calculated from the biological duplicates of the two experiments. The NBDC intensity and cell counts obtained from the NBDC experiments are available for re-analysis in Supplementary Data set 1, and raw flow cytometry plots are depicted in Supplementary Data set 2.

### Long read sequencing, assembly, and annotation

Cells were grown overnight in 500-mL shake flasks containing 100 mL liquid YPD medium at 30 °C in an orbital shaker at 200 rpm. After reaching stationary phase the cells were harvested for a total OD_660_ of 600 by centrifugation for 5 min at 4000 g. Genomic DNA of CBS6556 and NBRC1777 was isolated using the Qiagen genomic DNA 100/G kit (Qiagen, Hilden, Germany) according to the manufacturer’s instructions. MinION genomic libraries were prepared using the 1D Genomic DNA by ligation (SQK-LSK108) for CBS6556, and the 1D native barcoding Genomic DNA (EXP-NBD103 & LSK108) for NBRC1777 according to the manufacturer’s instructions with the exception of using 80% EtOH during the ‘End Repair/dA-tailing module’ step. Flow cell quality was tested by running the MinKNOW platform QC (Oxford Nanopore Technology, Oxford, UK). Flow cells were prepared by removing 20 μL buffer and subsequently primed with priming buffer. The DNA library was loaded dropwise into the flow cell for sequencing. The SQK-LSK108 library was sequenced on a R9 chemistry flow cell (FLO-MIN106) for 48 h. Base-calling was performed using Albacore (v2.3.1, Oxford Nanopore Technologies) for CBS6556, and for NBRC1777 with Guppy (v2.1.3, Oxford Nanopore Technologies) using dna_r9.4.1_450bps_flipflop.cfg. CBS6556 reads were assembled using Canu (v1.8)^69^, and NBRC1777 reads were assembled using Flye (v2.7.1-b1673)^70^. Assemblies were polished with Pilon (v1.18)^71^ using Illumina data available at the Sequence Read Archive under accessions SRX3637961 and SRX3541357. Both *de novo* genome assemblies were annotated using Funannotate (v1.7.1)^72^, trained and refined using *de novo* transcriptome assemblies (see below), adding functional annotation with Interproscan (v5.25-64.0)^73^.

### Illumina sequencing

Plasmids were sequenced on a MiniSeq (Illumina, San Diego, CA) platform. Library preparation was performed with Nextera XT DNA library preparation according to the manufacturer’s instructions (Illumina). The library preparation included the MiniSeq Mid Output kit (300 cycles) and the input & final DNA was quantified with the Qubit HS dsDNA kit (Life Technologies, Thermo Fisher Scientific). Nucleotide sequences were assembled with SPAdes^74^ and compared to the intended *in silico* DNA construct. For whole-genome sequencing, yeast cells were harvested from overnight cultures and DNA was isolated with the Qiagen genomic DNA 100/G kit (Qiagen) as described earlier. DNA quantity was measured with the QuBit BR dsDNA kit (Thermo Fisher Scientific). 300 bp paired-end libraries were prepared with the TruSeq DNA PCR-free library prep kit (Illumina) according to the manufacturer’s instructions. Short read whole-genome sequencing was performed on a MiSeq platform (Illumina).

### RNA isolation, sequencing and transcriptome analysis

Culture broth from chemostat cultures was directly sampled into liquid nitrogen to prevent mRNA turnover. The cell cultures were stored at −80 °C and processed within 10 days after sampling. After thawing on ice, cells were harvested by centrifugation. Total RNA was extracted by a 5 min heatshock at 65 °C with a mix of isoamyl alcohol, phenol and chloroform at a ratio of 125:24:1, respectively (Invitrogen). RNA was extracted from the organic phase with Tris-HCl and subsequently precipitated by the addition of 3 M Nac-acetate and 40 % (v/v) ethanol at −20 °C. Precipitated RNA was washed with ethanol, collected and after drying resuspended in RNAse free water. The quantity of total RNA was determined with a Qubit RNA BR assay kit (Thermo Fisher Scientific). RNA quality was determined by the RNA integrity number with RNA screen tape using a Tapestation (Agilent). RNA libraries were prepared with the TruSeq Stranded mRNA LT protocol (Illumina, #15031047) and subjected to paired-end sequencing (151 bp read length, NovaSeq Illumina) by Macrogen (Macrogen Europe, Amsterdam, the Netherlands).

Pooled RNAseq libraries were used to perform *de novo* transcriptome assembly using Trinity (v2.8.3)^75^ which was subsequently used as evidence for both CBS6556 and NBRC1777 genome annotations. RNAseq libraries were mapped into the CBS6556 genome assembly described above, using bowtie (v1.2.1.1)^76^ with parameters (-v 0 -k 10 --best -M 1) to allow no mismatches, select the best out of 10 possible alignments per read, and for reads having more than one possible alignment randomly report only one. Alignments were filtered and sorted using samtools (v1.3.1)^77^. Read counts were obtained with featureCounts (v1.6.0)^78^ using parameters (-B -C) to only count reads for which both pairs are aligned into the same chromosome.

Differential gene expression (DGE) analysis was performed using edgeR (v3.28.1)^79^. Genes with 0 read counts in all conditions were filtered out from the analysis, same as genes with less than 10 counts per million. Counts were normalized using the trimmed mean of M values (TMM) method^80^, and dispersion was estimated using generalized linear models. Differentially expressed genes were then calculated using a log ratio test adjusted with the Benjamini-Hochberg method. Absolute log2 fold-change values > 2, false discovery rate < 0.5, and P value < 0.05 were used as significance cutoffs.

Gene set analysis (GSA) based on gene ontology (GO) terms was used to get a functional interpretation of the DGE analysis. For this purpose, GO terms were first obtained for the *S. cerevisiae* CEN.PK113-7D (GCA_002571405.2) and *K. marxianus* CBS6556 genome annotations using Funannotate and Interproscan as described above. Afterwards, Funannotate compare was used to get (co)ortholog groups of genes generated with ProteinOrtho5^38^ using the following public genome annotations *S. cerevisiae* S288C (GCF_000146045.2), *K. marxianus* NBRC1777 (GCA_001417835.1), *K. marxianus* DMKU3-1042 (GCF_001417885.1), in addition to the new genome annotations generated here for *S. cerevisiae* CEN.PK113-7D, and *K. marxianus* CBS6556 and NBRC1777. Predicted GO terms for *S. cerevisiae* CEN.PK113-7D and *K. marxianus* CBS6556 were kept, and merged with those from corresponding (co)orthologs from *S. cerevisiae* S288C. Genes with term GO:0005840 (ribosome) were not considered for further analyses. GSA was then performed with Piano (v2.4.0)^40^. Gene set statistics were first calculated with the Stouffer, Wilcoxon rank-sum test, and reporter methods implemented in Piano. Afterwards, consensus results were derived by p-value and rank aggregation, considered significant if absolute Fold Change values > 1. ComplexHeatmap (v2.4.3)^81^ was used to draw GSA results into Fig. 2, highlighting differentially expressed genes found in a previous study^51^. DGE and GSA were performed using R (v4.0.2)^82^.

### Anaerobic growth experiments

Anaerobic shake-flask experiments were performed in a Bactron anaerobic workstation (BACTRON300-2, Sheldon Manufacturing, Cornelius, OR) at 30 °C. The gas atmosphere consisted of 85% N_2_, 10% CO_2_ and 5% H_2_ and was maintained anaerobic by a Pd catalyst. The catalyst was re-generated by heating till 160 °C every week and interchanged by placing it in the airlock whenever the pass-box was used. 50-mL Shake flasks were filled with 40 mL (80 % volumetric) media and placed on an orbital shaker (KS 130 basic, IKA, Staufen, Germany) set at 240 rpm inside the anaerobic chamber. Sterile growth media was placed inside the anaerobic chamber 24 h prior to inoculation to ensure complete removal of traces of oxygen.

The anaerobic growth ability of the yeast strains was tested on SMG-urea with 50 g·L^−1^ glucose at pH 6.0 with Tween 80 prepared as described earlier. The growth experiments were started from aerobic pre-cultures on SMG-urea media and the anaerobic shake flasks were inoculated at an OD_660_ of 0.2 (corresponding to an OD_600_ of 0.14). In order to minimize opening the anaerobic chamber, culture growth was monitored by optical density measurements inside the chamber using an Ultrospec 10 cell density meter (Biochrom, Cambridge, UK) at a 600 nm wavelength. When the optical density of culture no longer increased or decreased new shake-flask cultures were inoculated by serial transfer at an initial OD_600_ of 0.2.

### Laboratory evolution in low oxygen atmosphere

Adaptive laboratory evolution for strict anaerobic growth was performed in a Bactron anaerobic workstation (BACTRON BAC-X-2E, Sheldon Manufacturing) at 30 °C. 50-mL Shake flasks were filled with 40 mL SMG-urea with 50 g·L^−1^ glucose and including 420 mg·L^−1^ Tween 80. Subsequently the shake-flask media were inoculated with IMX2323 from glycerol cryo-stock at OD_660_ < 0.01 and thereafter placed inside the anaerobic chamber. Due to frequent opening of the pass-box and lack of catalyst inside the pass-box oxygen entry was more permissive. After the optical density of the cultures no longer increased, cultures were transferred to new media by 40-50x serial dilution. For IMS1111, IMS1112, IMS1113 three and for IMS1131, IMS1132, IMS1133 four serial transfers in shake-flask media were performed after which single colony isolates were made by plating on YPD agar media with hygromycin antibiotic at 30 °C aerobically. Single colony isolates were subsequently restreaked sequentially for three times on the same media before the isolates were propagated in SM glucose media and glycerol cryo stocked.

To determine if an oxygen-limited pre-culture was required for the strict anaerobic growth of IMX2323 strain a cross-validation experiment was performed. In parallel, yeast strains were cultivated in 50-mL shake-flask cultures with SMG-urea with 50 g·L^−1^ glucose at pH 6.0 with Tween 80 in both the Bactron anaerobic workstation (BACTRON BAC-X-2E, Sheldon Manufacturing) with low levels of oxygen-contamination, and in the Bactron anaerobic workstation (BACTRON300-2, Sheldon Manufacturing) with strict control of oxygen-contamination. After stagnation of growth was observed in the second serial transfer of the shake-flask cultures a 1.5 mL sample of each culture was taken, sealed, and used to inoculate fresh-media in the other Bactron anaerobic workstation. Simultaneously, the original culture was used to inoculate fresh media in the same Bactron anaerobic workstation, thereby resulting in 4 parallel cultures of each strain of which halve were derived from the other Bactron anaerobic workstation.

### Laboratory evolution in sequential batch reactors

Laboratory evolution for selection of fast growth at high temperatures was performed in 400-mL MultiFors (Infors Benelux, Velp, the Netherlands) bioreactors with a working volume of 100 mL for the strain IMS1111 on SMG 20 g·L^−1^ glucose media with Tween 80 in triplicate. Anaerobic conditions were created and maintained by continuous aeration of the cultures with 50 mL·min^−1^ (0.5 vvm) N_2_ gas and continuous aeration of the media vessels with N_2_ gas. The pH was set at 5.0 and maintained by the continuous addition of sterile 2 M KOH. Growth was monitored by analysis of the CO_2_ in the bioreactor off-gas and a new empty-refill cycle was initiated when the batch time had at least elapsed 15 hours and the CO_2_ signal dropped to 70% of the maximum reached in each batch. The dilution factor of each empty-refill cycle was 14.3-fold (100 mL working volume, 7 mL residual volume). The first batch fermentation was performed at 30 °C after which in the second batch the temperature was increased to 42 °C and maintained at for 18 consecutive sequential batches. After the 18 batch cycle at 42 °C the culture temperature was again increased to 45 °C and maintained subsequently. Growth rate was calculated based on the CO_2_ production as measured by the CO_2_ fraction in the culture off-gas in essence as described previously^83^. In short, the CO_2_ fraction in the off-gas was converted to a CO_2_ evolution rate of mmol per hour and subsequently summed over time for each cycle. The corresponding cumulative CO_2_ profile was transformed to natural log after which the stepwise slope of the log transformed data was calculated. Subsequently an iterative exclusion of datapoints of the stepwise slope of the log transformed cumulative CO_2_ profile was performed with exclusion criteria of more than one standard deviation below the mean.

### Variant calling

DNA sequencing reads were aligned into the NBRC1777 described above including an additional sequence with *TtSTC1* construct, and used to detect sequence variants using a method previously reported^84^. Briefly, reads were aligned using BWA (v0.7.15-r1142-dirty)^85^, alignments were processed using samtools (v1.3.1)^77^ and Picard tools (v2.20.2-SNAPSHOT) (http://broadinstitute.github.io/picard), and variants were then called using the Genome Analysis Toolkit (v3.8-1-0-gf15c1c3ef)^86^ HaplotypeCaller in DISCOVERY and GVCF modes. Variants were only called at sites with minimum variant confidence normalized by unfiltered depth of variant samples (QD) of 20, read depth (DP) ≥ 5, and genotype quality (GQ) > 20, excluding a 7.1 kb region in chromosome 5 containing rDNA. Variants were annotated using the genome annotation described above, including the *TtSTC1* construct, with SnpEff (v5.0)^87^ and VCFannotator (http://vcfannotator.sourceforge.net).

### Statistics

Statistical test performed are given as two sided with unequal variance t-test unless specifically stated otherwise. We denote technical replicates as measurements derived from a single cell culture. Biological replicates are measurements originating from independent cell cultures. Independent experiments are two experiments identical in set-up separated by the difference in execution days. If possible variance from independent experiments with identical setup were pooled together, but independent experiments from time-course experiments (anaerobic growth studies) are reported separately. *p*-values were corrected for multiple-hypothesis testing which is specifically reported each time. No data was excluded based on the resulting data out-come.

### Data availability

Data supporting the findings of this work are available within the paper and source data for all figures in this study are available at the www.data.4TU.nl repository with the doi:10.4121/13265552.

The raw RNA-sequencing data that supports the findings of this study are available from the Genome Expression Omnibus (GEO) website (https://www.ncbi.nlm.nih.gov/geo/) with number GSE164344. Whole-genome sequencing data of the CBS6556, NBRC1777 and evolved strains were deposited at NCBI (https://www.ncbi.nlm.nih.gov/) under BioProject accession number PRJNA679749.

### Code availability

The code that were used to generate the results obtained in this study are archived in a Gitlab repository (https://gitlab.tudelft.nl/rortizmerino/kmar_anaerobic).

## Author’s contributions

WD and JTP designed the study and wrote the manuscript. WD performed molecular cloning, bioreactor cultivation experiment, transcriptome analysis and sterol-uptake experiments. JB contributed to bioreactor cultivation experiments and molecular cloning. FW contributed to the molecular cloning and sterol-uptake experiments. AK and CM contributed to bioreactor experiments and transcriptome studies. PdlT performed plasmid and genome sequencing. RO contributed to transcriptome analysis and performed sequence annotation and assembly.

## Acknowledgements

We thank Mark Bisschops and Hannes Jürgens for fruitful discussions. We thank Erik de Hulster for fermentation support and Marcel van den Broek for input on the bioinformatics analyses.

## Description of Additional Supplementary Files

**Supplementary Data Set 1 Overview of flow cytometry samples with meta-data.** Meta-data Table of file names, frequency of cells compared to parent, number of cells in each group, strain name, time point of fluorescence measurement after 4 hours (1) or 23 hours (2), staining of cells with propidium-iodide (PI) with value (PI) or without PI staining (-), staining of cells with Tween 80 NBD-cholesterol (TN) or with Tween 80 only (T), with species names abbreviated *K. marxianus* (Km) or *S. cerevisiae* (Sc).

[Example picture of file FlowCyto_Table.xlsx]

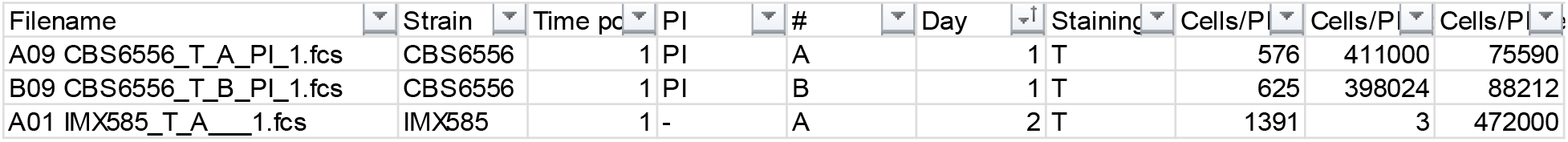

**Supplementary Data set 2 Flow cytometry non-gated data of FL3-A versus FL1-A of all samples.**

Flow cytometry data of showing fluorescent NBDC uptake by *K. marxianus*, *S. cerevisiae* strains with for each sample the intensity of counts (pseudo-colored) for 533/30 nm (FL1) for NBDC and > 670 nm (FL3) for PI.

[Example of first row of FlowCyto_FL1_FL3.pdf]

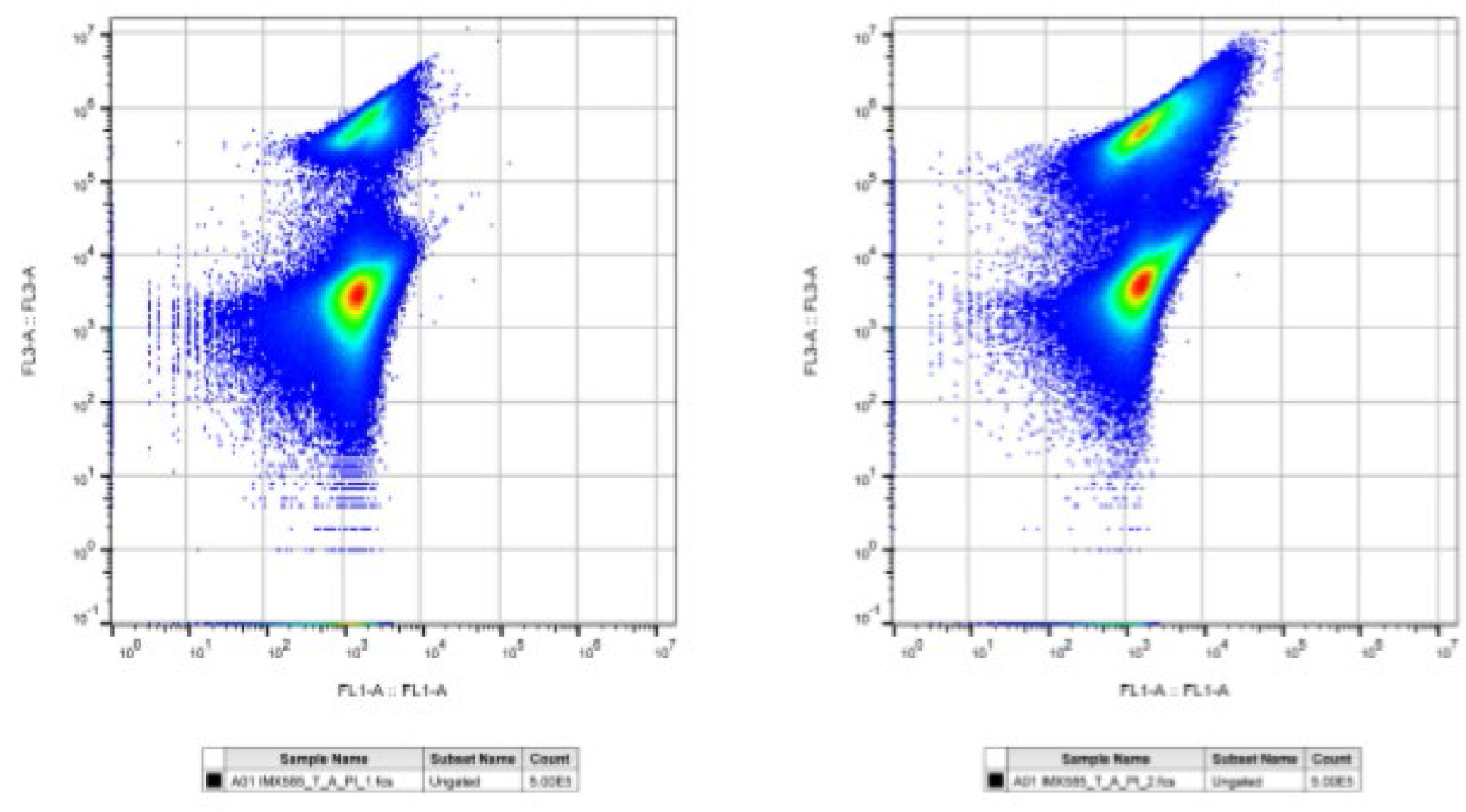

## Supplemental material for

**Supplementary Fig. 1.**
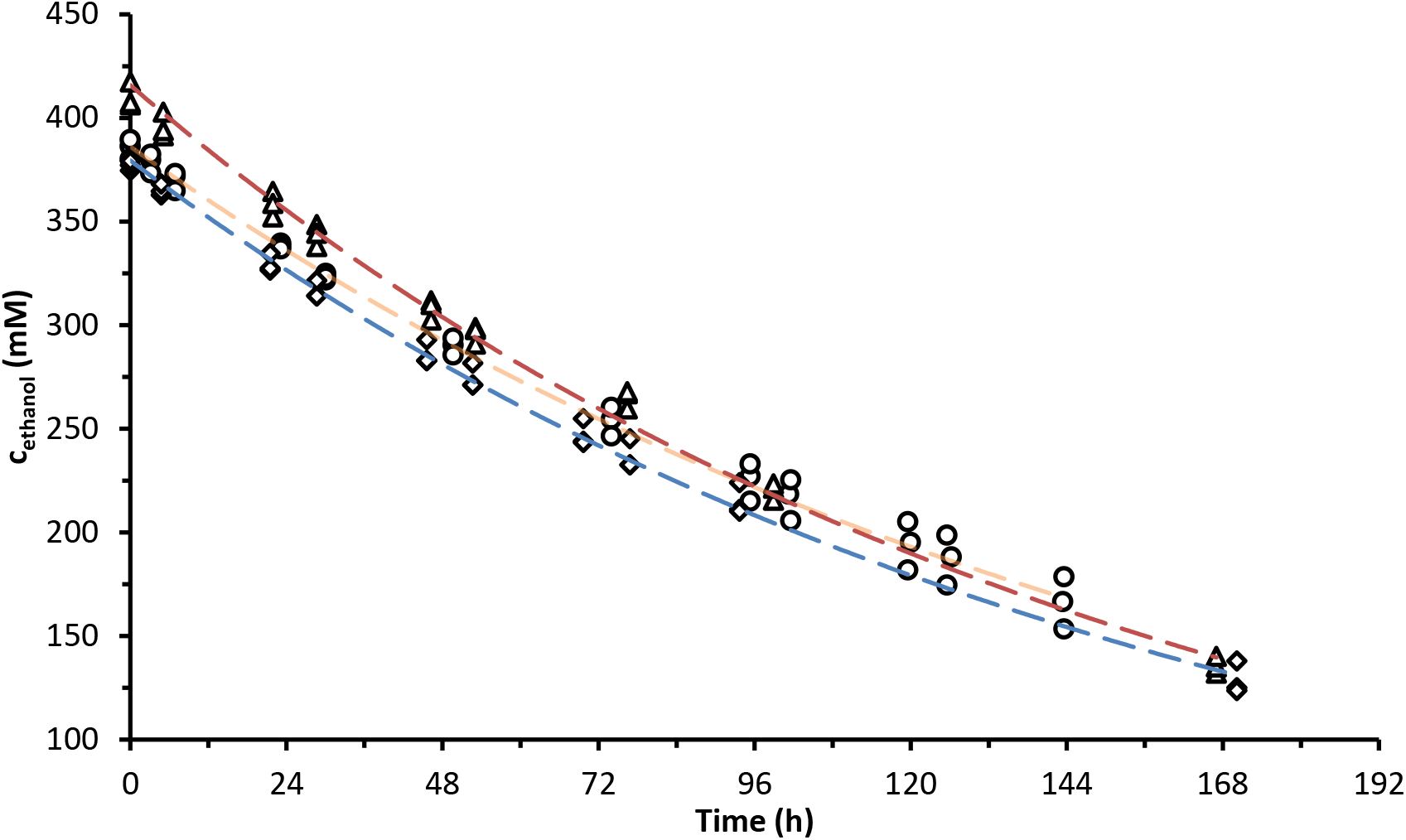
Ethanol evaporation rate. Ethanol concentration over time with reactor volume of 1200 mL SM glucose urea media maintained at 30 °C, stirred with 800 rpm and aerated with a volumetric gas flow rate of 500 mL·min^−1^. The reactor off-gas was cooled by passing through a condenser cooled at 2 °C. Circles and orange line represent the condition with sparge aeration and Tween 80 (T) media supplementation, diamonds and blue line head-space aeration with Tween 80, triangle and red line represent head space aeration and Tween 80 omission. Data represent mean with standard deviation from three independent reactor experiments.

| AGF | Aeration type | Ethanol evaporation (mmol·h<sup>-1</sup>) |
| --- | --- | --- |
| T | Sparge | 0.00578 ± 0.00062 |
| T | Head-space | 0.00625 ± 0.00032 |
|  | Head-space | 0.00653 ± 0.00020 |

**Supplementary Fig. 2.**
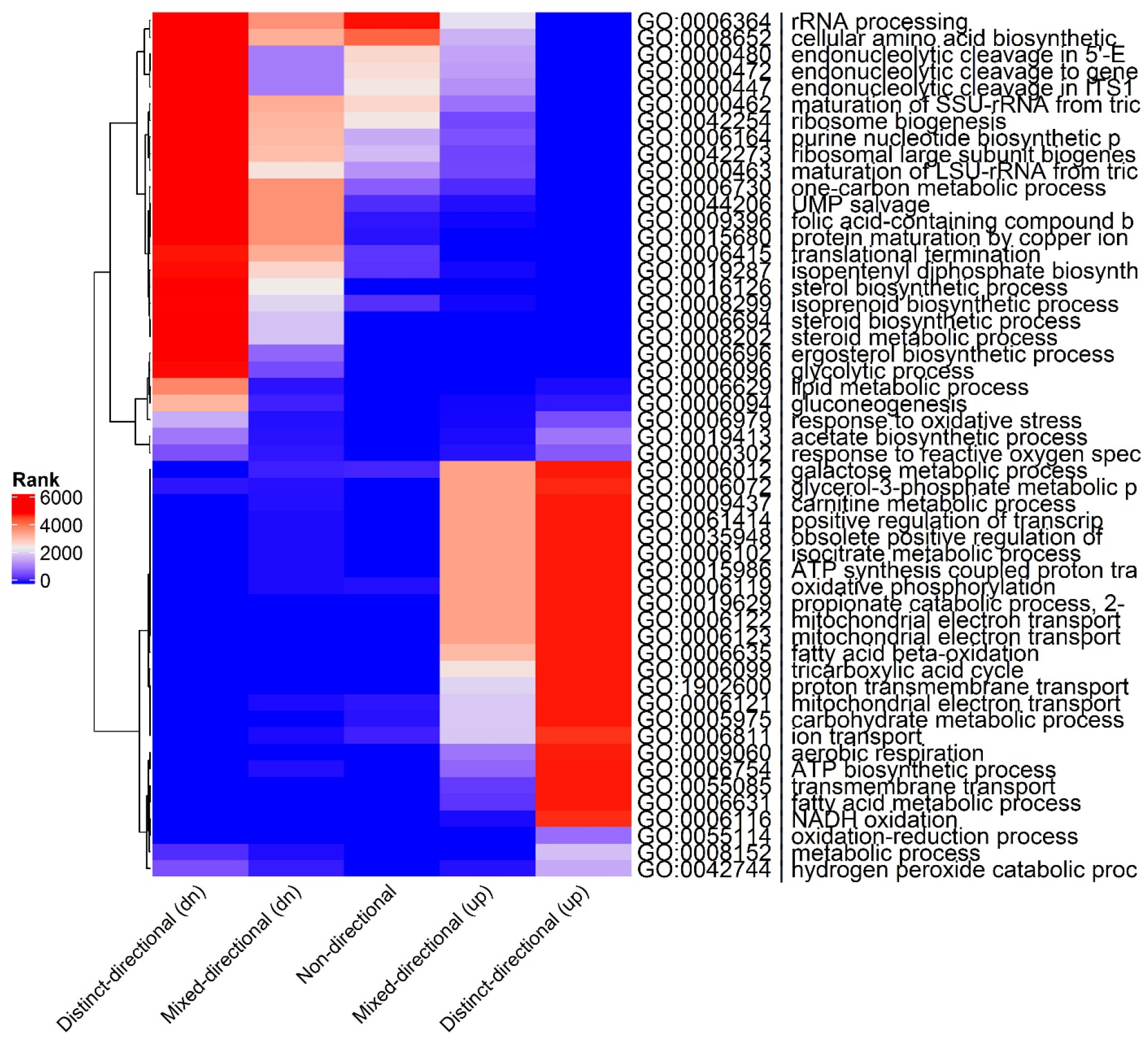
Consensus biological process GO term enrichment for *K. marxianus* contrast 31. GO terms are clustered according to their rank. See legend of Fig. 2 for experimental details.

**Supplementary Fig. 3.**
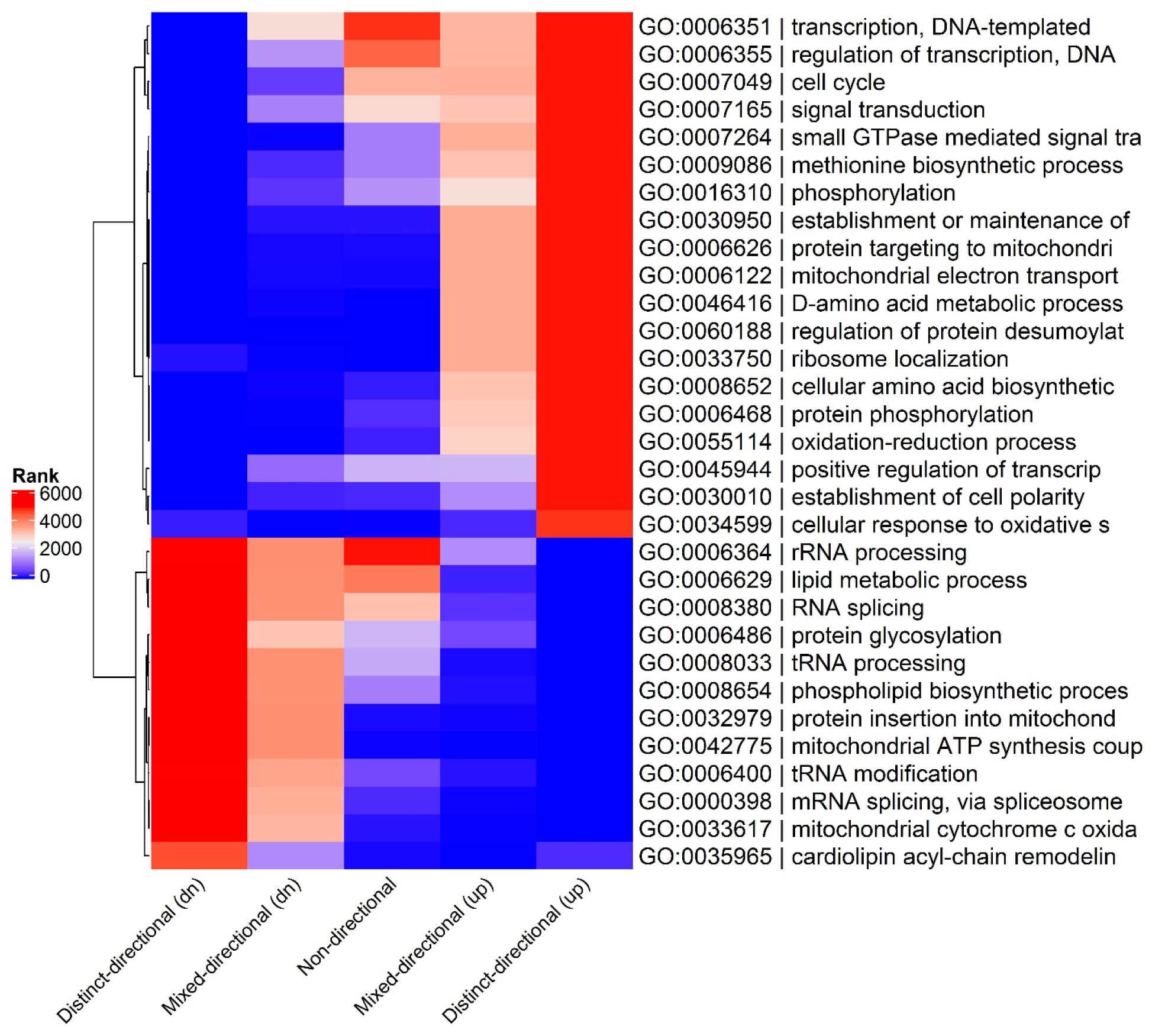
Consensus biological process GO term enrichment for *K. marxianus* contrast 43. GO terms are clustered according to their rank. See legend of Fig. 2 for experimental details.

**Supplementary Fig. 4.**
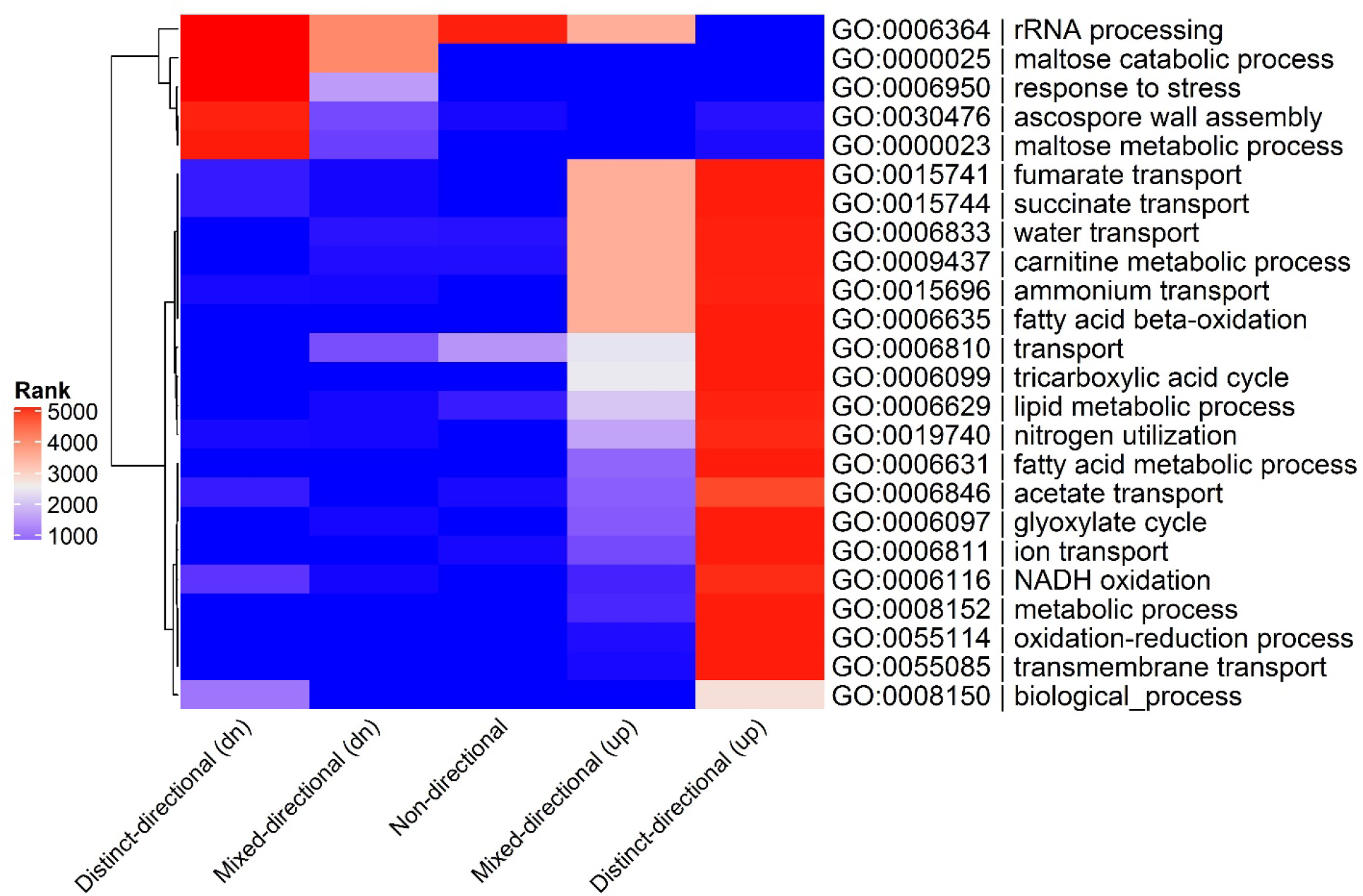
Consensus biological process GO term enrichment for *S. cerevisiae* contrast 31. GO terms are clustered according to their rank. See legend of Fig. 2 for experimental details.

**Supplementary Fig. 5.**
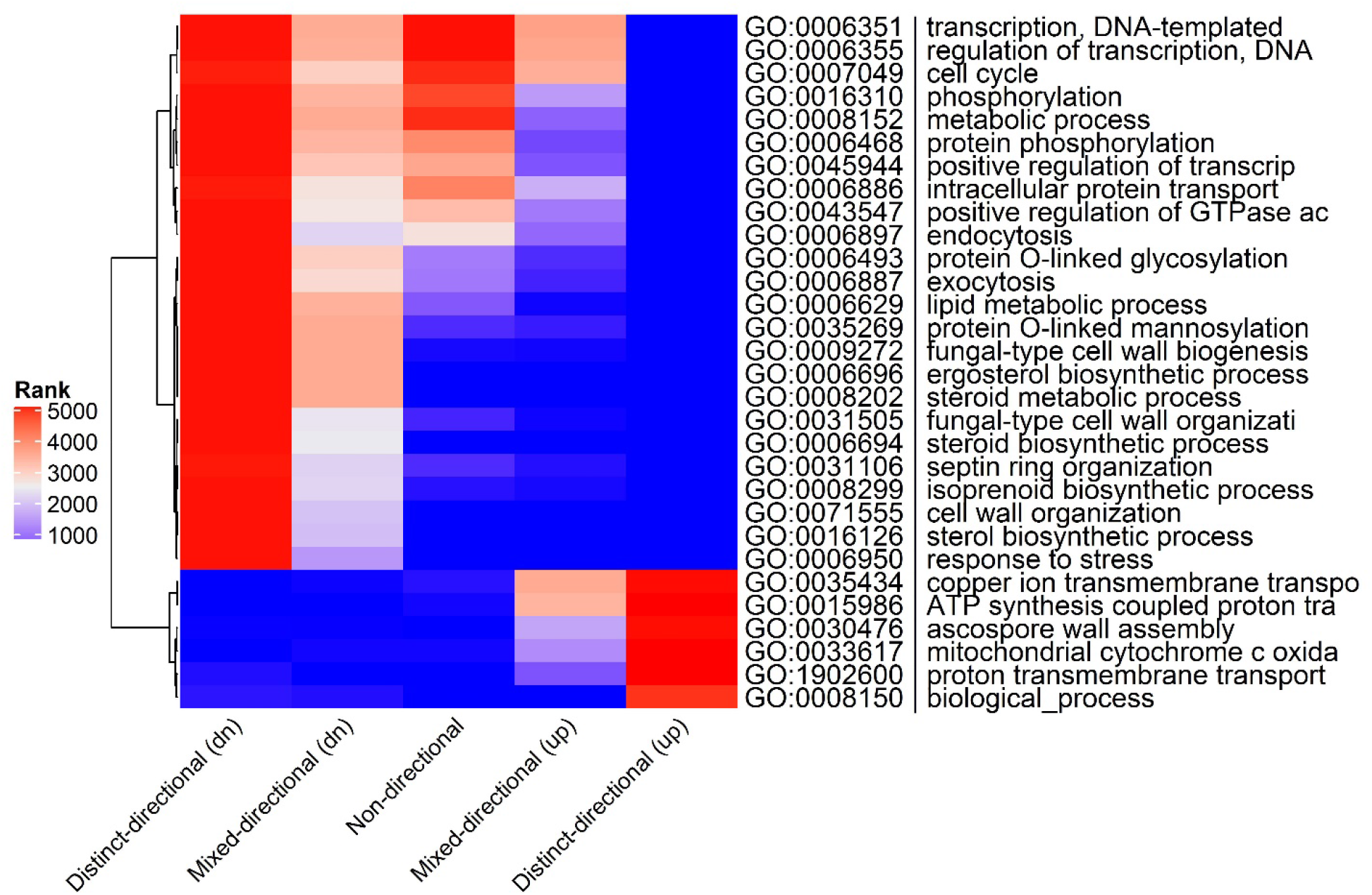
Consensus biological process GO term enrichment for *S. cerevisiae* contrast 43. GO terms are clustered according to their rank. See legend of Fig. 2 for experimental details.

**Supplementary Fig. 6.**
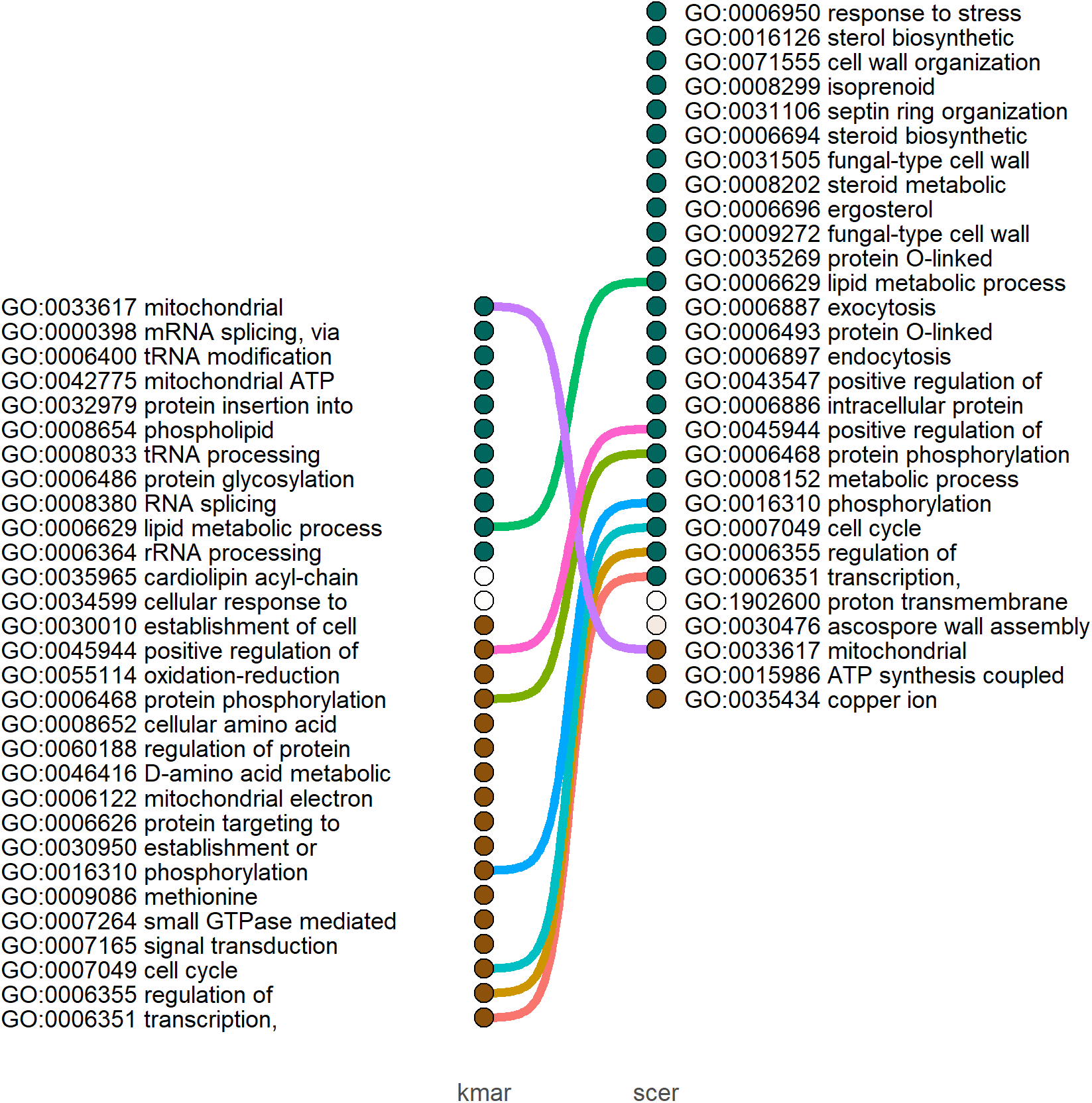
GO term enrichment comparison of biological process of *K. marxianus* (kmar) to *S. cerevisiae* (scer) of contrast 43. GO terms were annotated with the color of distinct directionality (up (blue) down (brown)) and the color intensity was determined by the magnitude of the inverse rank. GO terms with significant mixed-directionality or non-directionality, as having no pronounced distinct directionality, are colored white. Shared GO terms between *K. marxianus* and *S. cerevisiae* are connected by a line.

**Supplementary Fig. 7.**
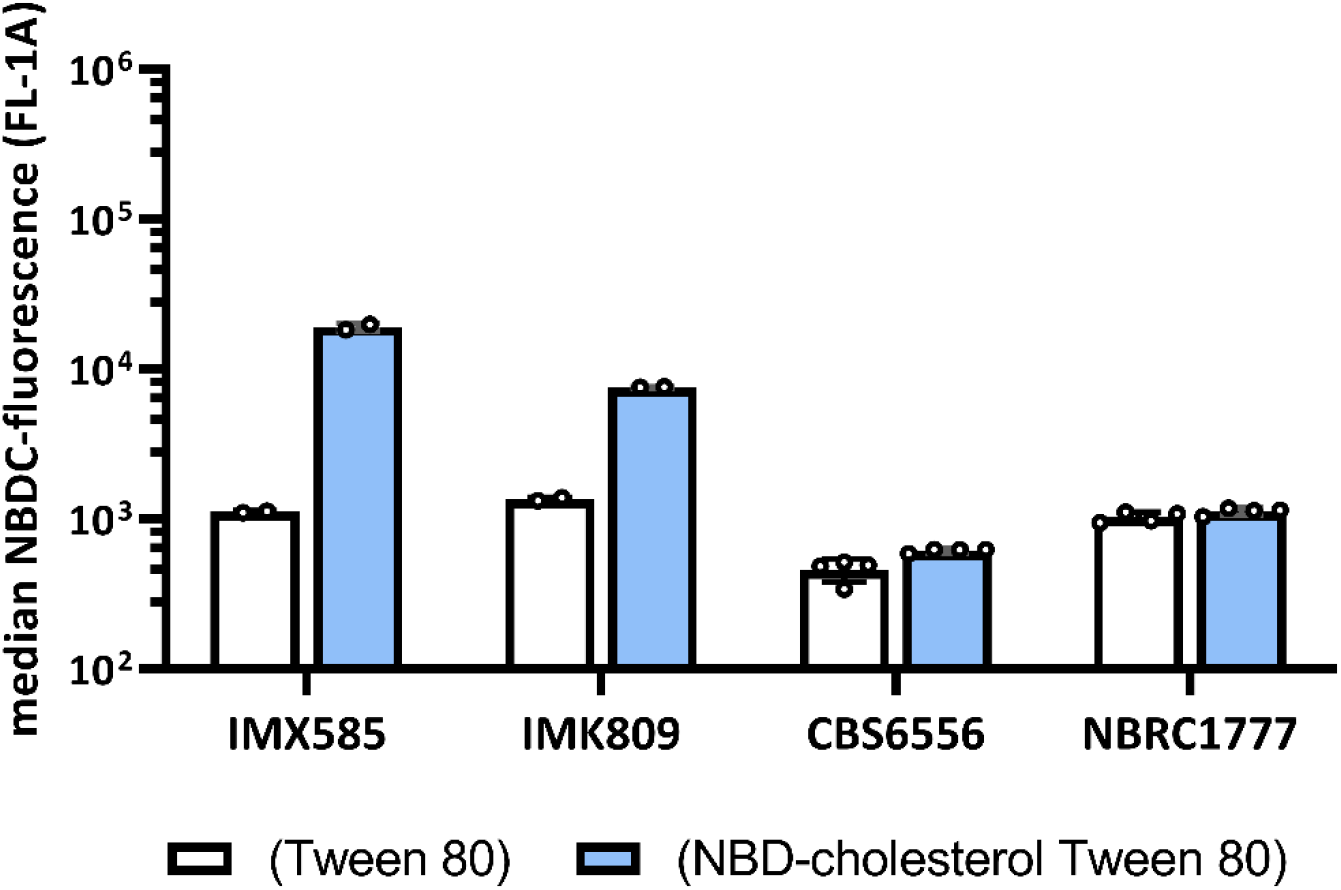
Uptake of the fluorescent sterol derivative NBDC by *S. cerevisiae* and *K. marxianus* strains after 23 h staining. Flow cytometry data of Fig. 4 with prolonged staining after pulse-addition of NBD-cholesterol to the shake-flask cultures for 23 h. Bar charts of the median and pooled standard deviation of the NBD-cholesterol fluorescence intensity of PI-negative cells with pooled variance from the biological replicate cultures. See legend Fig. 4 for experimental details.

**Supplementary Fig. 8.**
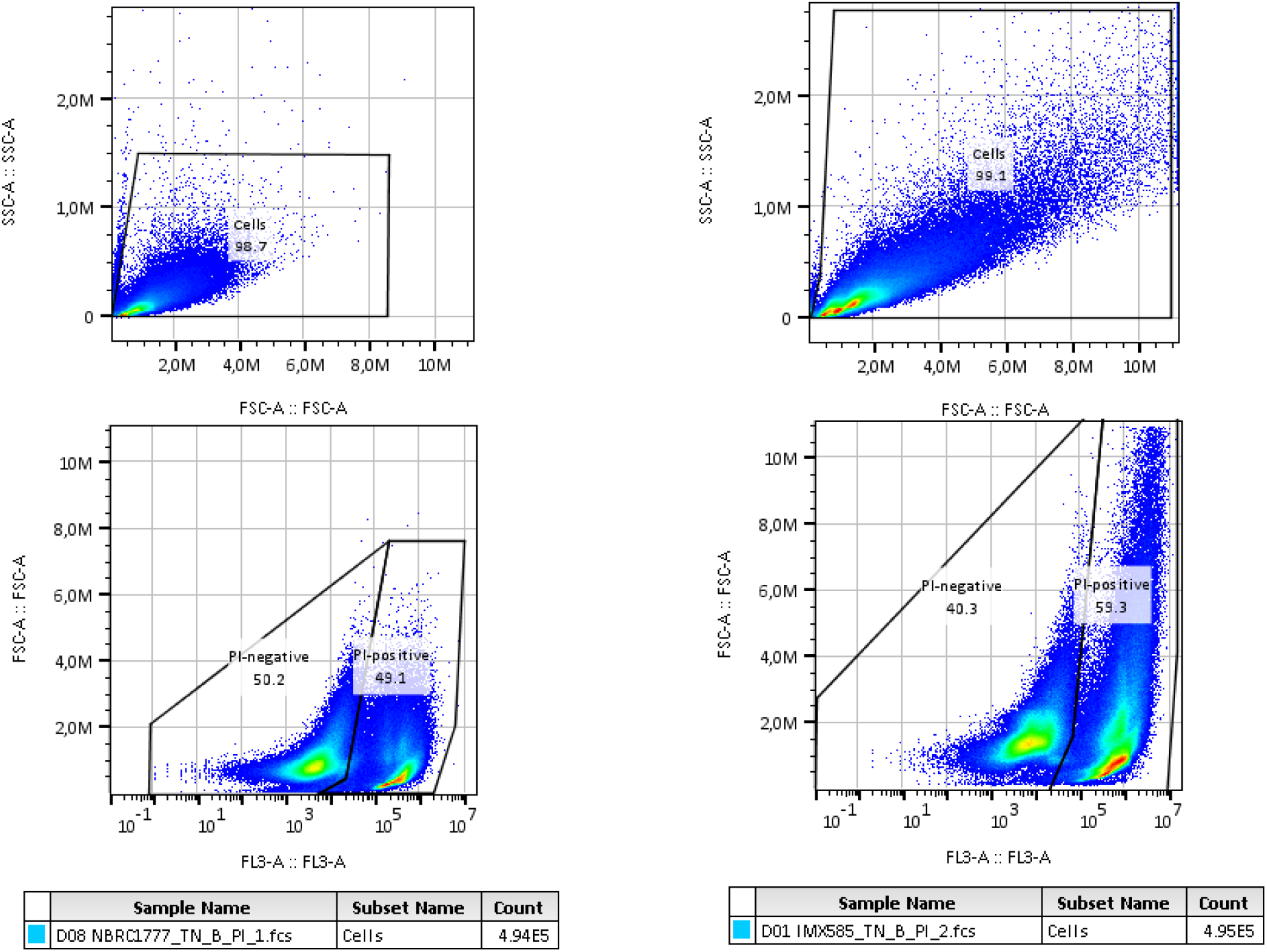
Flow cytometry gating strategy of both *K. marxianus* (left panel) and *S. cerevisiae* (right panel) samples. Gates were set per one species for all samples independent of NBDC staining. Density of events were calculated by FlowJo software and represented in pseudo-color (blue low density, red high-density). The gate between PI-negative and PI-positive was inside the “Cells” gated-population.

**Supplementary Fig. 9.**
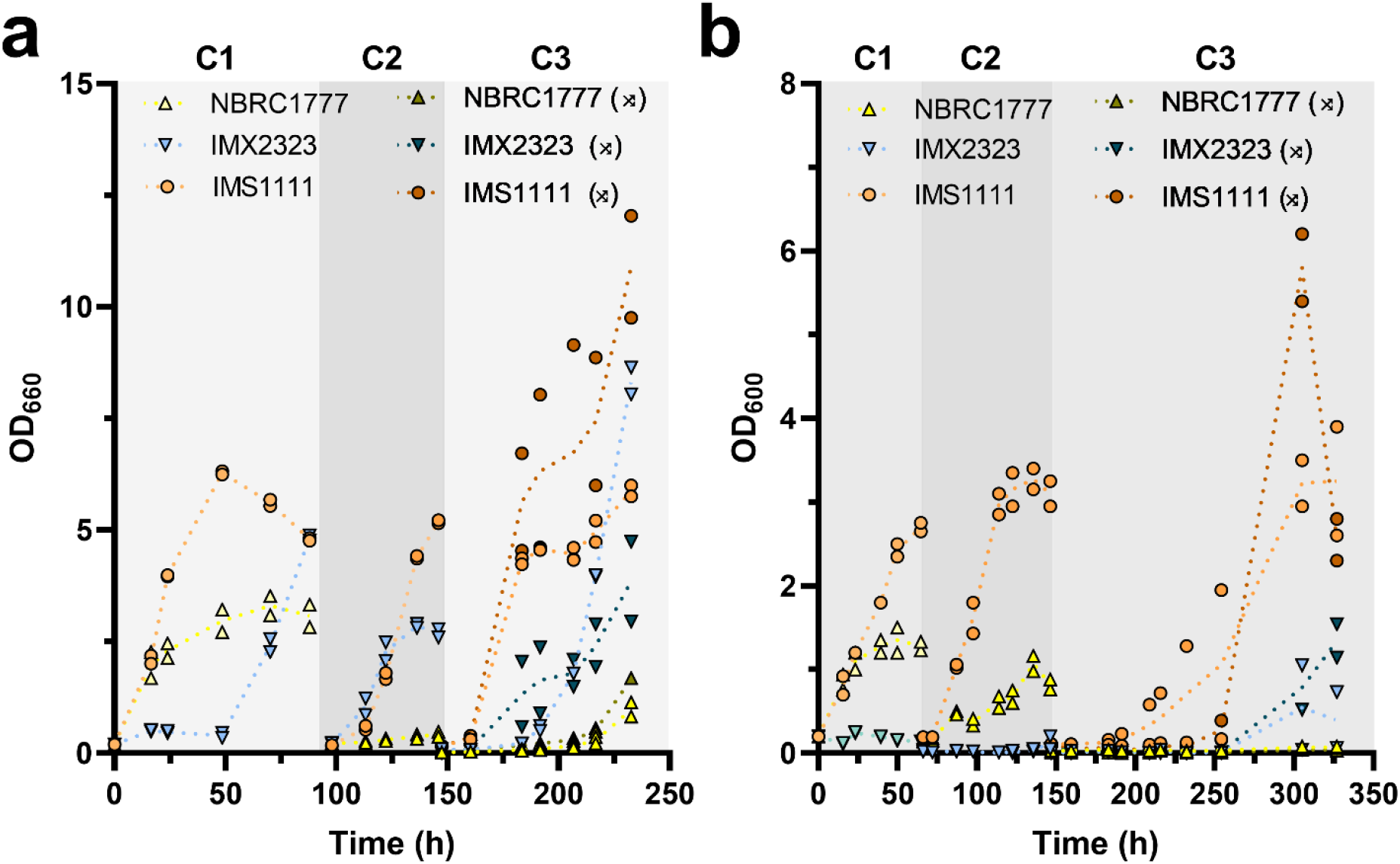
Cross-validation of oxygen-limited and anaerobic growth of *K. marxianus* IMX2323. Strains were grown in shake-flask cultures in an oxygen-limited (**a**) and strict anaerobic environment (**b**). To perform cross-validation between the two parallel running experiments, 1.5 mL aliquot of each culture was sealed and transferred quickly between anaerobic chambers and used to inoculate two shake-flask cultures, represented with crossed-arrows (⤮). The cultures from the strain NBRC1777 (⤮) in the third transfer (C3) in the strict anaerobic environment (**b**) were hence inoculated from an aliquot of the cultures of NBRC1777 (C2) grown in oxygen-limited environment (**a**). This resulted in a serial transfer of 26.7 times dilution from transfer C2 to C3. Aerobic grown pre-cultures were used to inoculate the first anaerobic culture on SMG-urea containing 50 g·L^−1^ glucose and Tween 80. Data depicted are of each replicate culture (points) and the mean (dotted line) from independent biological duplicate cultures, serial transfers cultures are represented with the number of respective transfer (C1-3)

**Supplementary Fig. 10.**
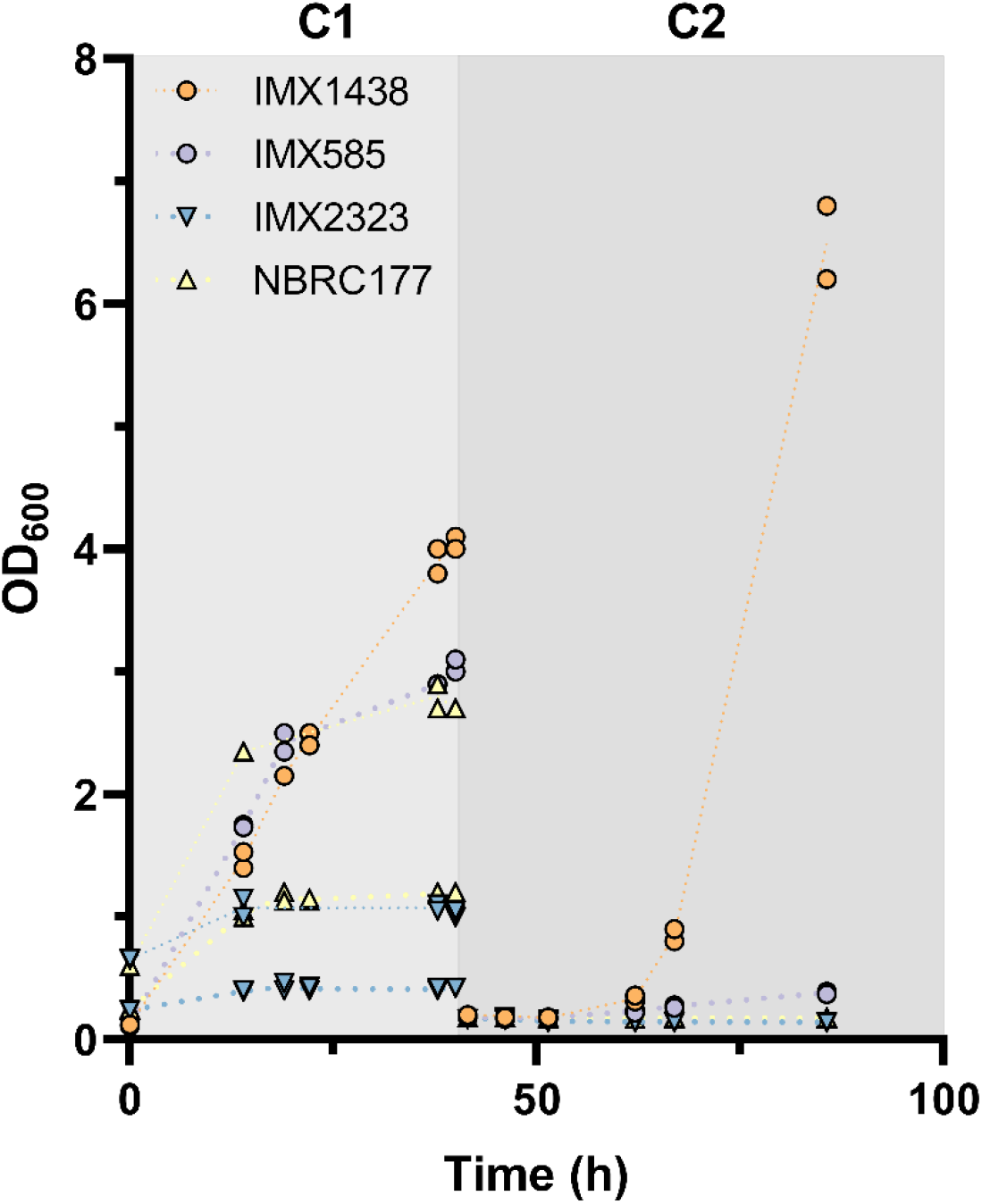
Sterol-independent anaerobic growth of *S. cerevisiae* IMX585 (reference), IMX1438 (*TtSTC1*), *K. marxianus* NBRC1777 (reference) and IMX2323 (*TtSTC1*). Aerobic grown pre-cultures were used to inoculate shake-flask cultures with SMG-urea containing 50 g·L^−1^ glucose and Tween 80 in a strict anaerobic environment at an OD_600_ of 0.1 for all strains, and both at OD_600_ of 0.1 and 0.6 for NBRC1777 and IMX2323. Data depicted are of each replicate culture (points) and the mean (dotted line) from independent biological duplicate cultures, serial transfers cultures are represented with the number of respective transfer (C1-2).

**Supplementary Fig. 11.**
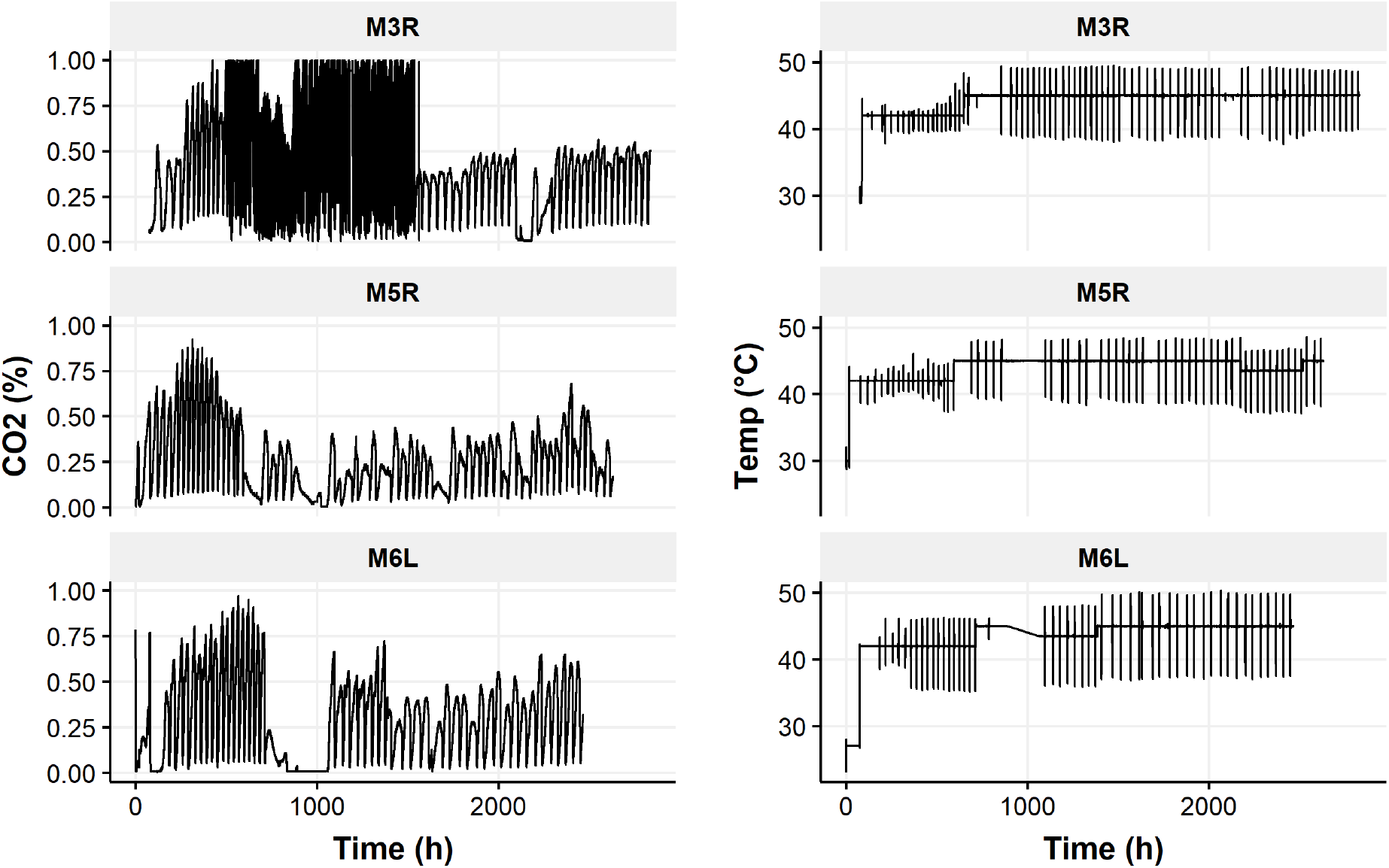
CO_2_ fraction in the off-gas of *K. marxianus* IMS1111. Production of CO_2_ as measured by the fraction of CO_2_ in the off-gas of the individual bioreactor cultivations of the *K. marxianus* strain IMS1111 on SMG media pH 5.0 with 20 g·L^−1^ glucose, 420 mg·L^−1^ Tween 80 over time (Left panels). The temperature profile was incrementally increased at the beginning of a new batch cycle (right panels). After 430 h the performance of the off-gas analyzer of replicate M3R deteriorated.

**Supplementary Table 1.**
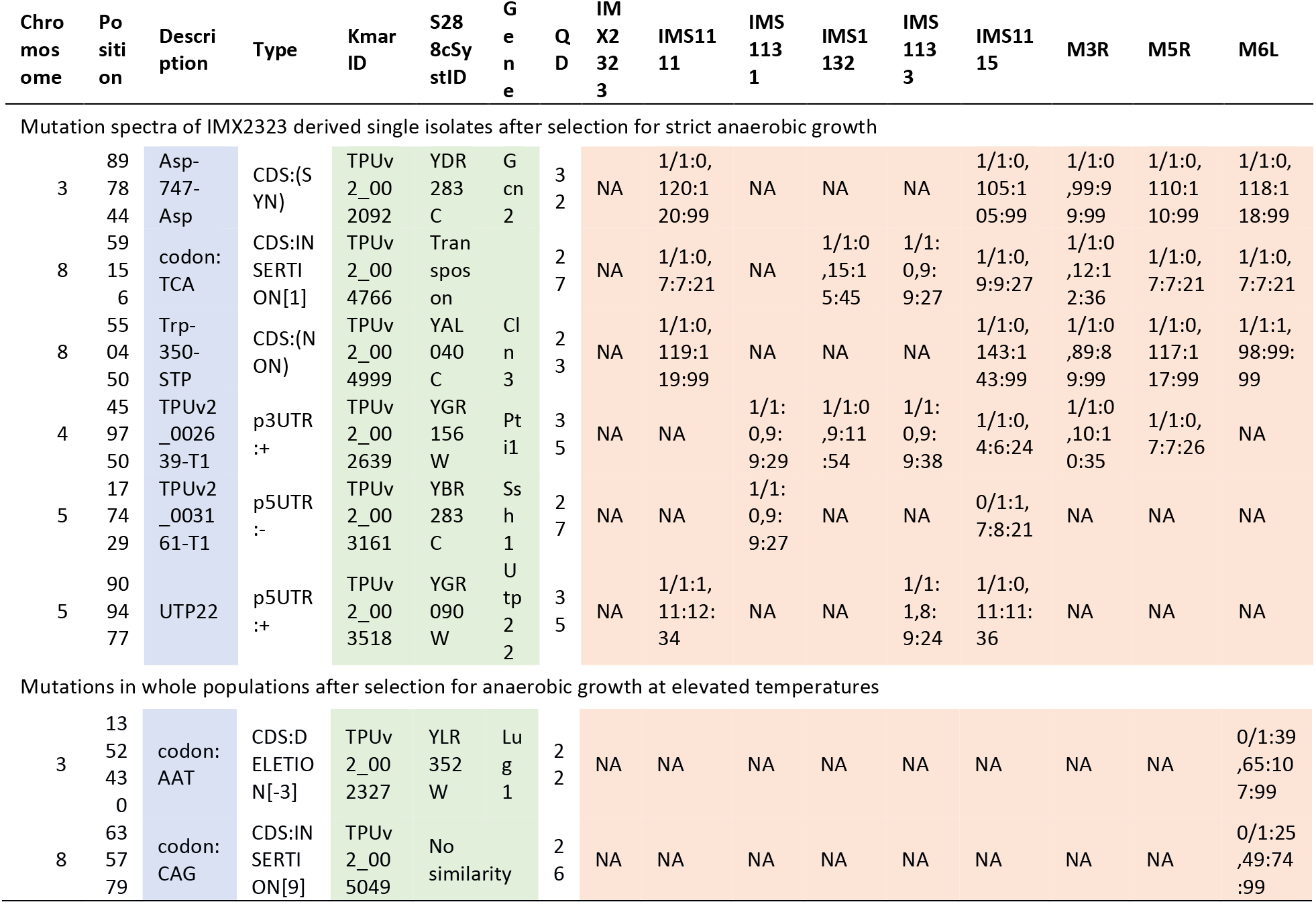
Mutations identified by whole-genome sequencing in comparison to the reference *K. marxianus* strain IMX2323. Overview of mutations detected in the strains after selected for strict anaerobic growth IMS1111, IMS1131, IMS1132, IMS1133 compared to the *TtSTC1* engineered strain (IMX2323). Resequencing of IMS1111 after 4 transfers in strict anaerobic conditions is for clarity referred with the strain name IMS1115. Overview of mutations of the bioreactor populations after prolonged selection for anaerobic growth at elevated temperatures, represented by the bioreactor replicates (M3R, M5R, and M6L). Mutations in coding regions are annotated as synonymous (SYN), non-synonymous (NSY), insertion or deletions. Mutations in non-coding regions are reported with the identifier of the neighboring gene, directionality and strand (+/−). For *K. marxianus* genes, corresponding *S. cerevisiae* orthologs with the S288C identifier are listed if applicable. QD refers to quality by depth calculated by GATK and genotyping overviews are given per strain using the GATK fields GT: 1/1 for homozygous alternative, 1/0 for heterozygous, AD: allelic depth (number of reads per reference and alternative alleles called), DP: approximate read depth at the corresponding genomic position, and GQ: genotype quality. NA indicates variants were not called in that position in the corresponding strain.

